# Billi: Accurate and Scalable Bubble Detection in Pangenome Graphs

**DOI:** 10.1101/2025.11.21.689636

**Authors:** Shreeharsha G Bhat, Daanish Mahajan, Chirag Jain

**Author notes:** Should be regarded as joint first authors (Shreeharsha G Bhat, Daanish Mahajan).

## Abstract

A key application of pangenome graphs is the characterization of small and large genomic variants by enumerating bubble-like structures in the graph. Several formalizations of bubbles have been proposed. Superbubbles and ultrabubbles are two such formalizations, but both exclude cyclic subgraphs. Representing cyclic structures is essential for capturing complex genomic variation, such as inversions. Other definitions that accommodate cycles do not guarantee that overlapping bubbles are strictly nested. This makes it difficult to organize variants hierarchically and distinguish complex variants from their constituent subvariants.

We introduce two formalizations, called *panbubble* and *hairpin*, that accommodate cyclic subgraphs. We prove that both satisfy the strict nesting property. In our theoretical analysis and algorithm design, we leverage an auxiliary undirected graph representation derived from the bidirected pangenome graph, enabling us to exploit established graph-theoretic algorithms and properties. We then present an exact *O*(|*V*|^2^ (|*V*| + |*E*|))-time algorithm for detecting all panbubbles and hairpins in a bidirected graph *G* = (*V, E*). In addition, we propose a heuristic that produces the same output as the exact algorithm on all tested graphs and scales to both the first and second releases of the Human Pangenome Reference Consortium. We implemented our algorithms in the tool `Billi.` At a known recurrent inversion locus on the human Y chromosome, Billi recovers the inversion together with its nested variants, whereas existing bubble-detection methods miss part of the variation or fail to correctly represent the nesting relationships among variants.

**Source code:** https://github.com/at-cg/billi

## 1 Introduction

Pangenome graphs provide a flexible framework for representing complex structural variations, including cases where smaller variations are nested within larger ones [2,4,17,24,34,35]. Unlike conventional variant calling methods, variant discovery in pangenome graphs is not constrained to the coordinate system of a single reference genome [1]. A key computational challenge in this setting is the identification of *bubbles*, which are substructures in the graph that likely correspond to genomic variation. Informally, a bubble is a region of the graph where multiple paths diverge from a common source vertex and subsequently reconverge at a common sink vertex [15]. Bubbles and efficient algorithms for their detection were first studied in the context of genome assembly [3,12,26,36]. However, these studies assume a directed graph representation. In contrast, most pangenome graph construction methods [11,14,18,19] use *bidirected graphs* to represent the double-stranded nature of DNA.

Efforts to generalize the concept of bubbles to bidirected graphs [8,19,23,27] build upon the notion of the *super-bubble* [26]. Superbubbles are an important class of subgraphs in directed graphs, originally introduced to simplify genome assembly graphs by identifying substructures that arise from sequencing errors, heterozygous variants, and near-identical repeats [38]. Onodera *et al*. [26] gave the conditions under which a pair of distinct vertices *s* and *t* can serve as the source and sink of a superbubble. Specifically, (i) the enclosed subgraph *B* is acyclic, (ii) no edge enters *B*\{*s*} from outside *B*, and (iii) no edge leaves *B* \{*t*} to a vertex outside *B*. Superbubbles exhibit elegant structural properties. For instance, the interior subgraphs of two superbubbles are either disjoint or strictly nested, and every vertex within a superbubble lies on some source-to-sink path. *Weak superbubbles* relax the superbubble definition by permitting the otherwise forbidden edge from *t* to *s* [12].

For bidirected graphs, the generalizations include *ultrabubbles* [27], *snarls* [27], *bibubbles* [19], and *flubbles* [23]. In pangenome graphs, cyclic subgraphs are of particular interest because genomic rearrangements such as inversions and duplications can give rise to cyclic structures (Figure 1B). Snarls, bibubbles, and flubbles relax the acyclicity constraint and can therefore represent such structures, but none fully preserves the properties of superbubbles. Snarls and flubbles may contain vertices that are not part of any source-to-sink walk (Figure 1C). Pairs of snarls, bibubbles, or flubbles may overlap without being strictly nested (Figure 1D). Such overlaps can introduce ambiguity in delineating the region corresponding to a given variant. Section 2.6 in [19] claims that overlapping bibubbles exhibit strict nesting without a proof. However, the counterexamples in Figure 1D and Supplementary Figure S1 suggest that this claim does not hold in general. For snarls, the existing remedy is to use heuristics and select a non-overlapping subset, termed *compatible snarls* [27].

**Fig. 1:**
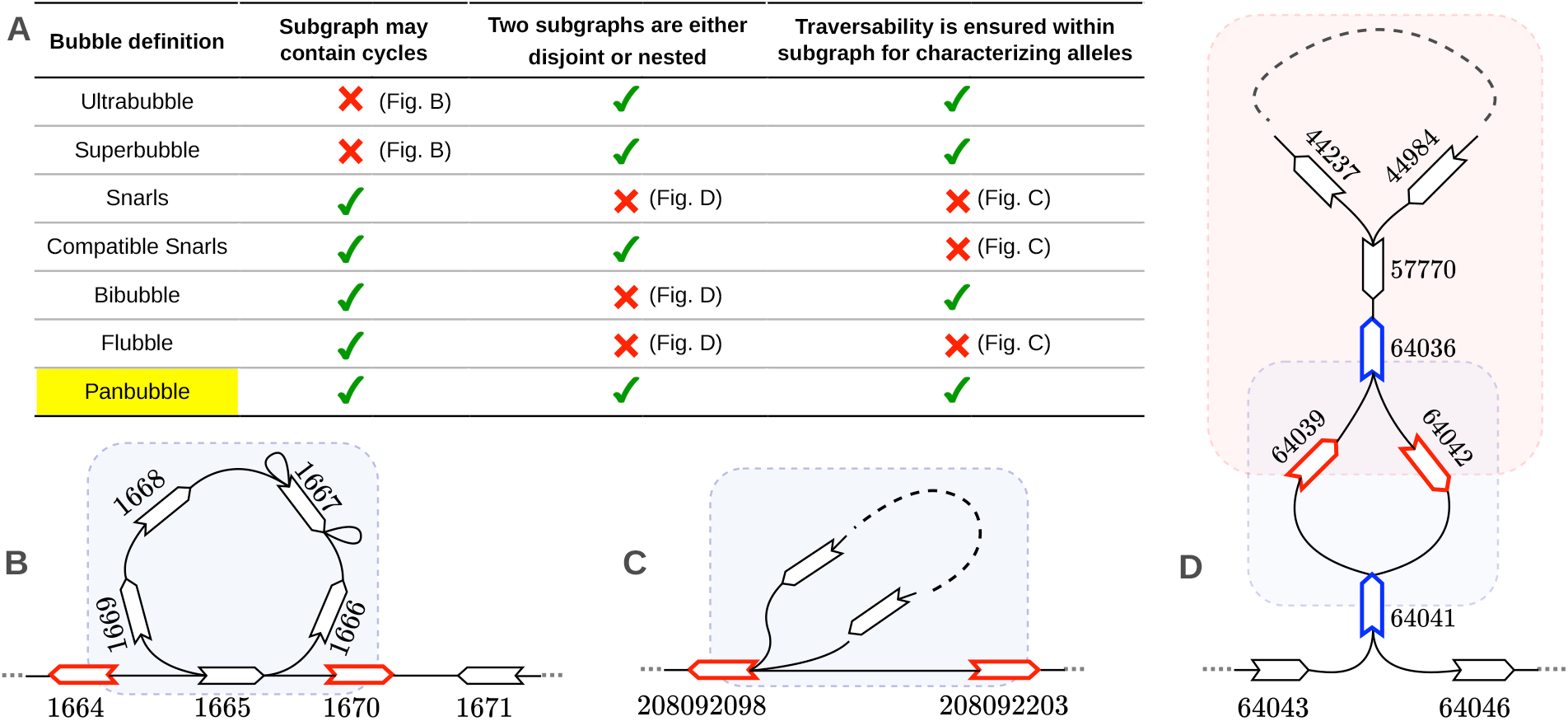
**(A)** Summary of the structural properties of different bubble definitions. Although superbubbles were originally defined for directed graphs [26], we include them here because they can also be applied to a bidirected graph [8]. **(B-D)** Bidirected subgraphs extracted from publicly-available pangenome graphs: `MUC4`, `HPRCv2.0-chrX`, and `HPRCv1.1-chrY`, respectively [6,20,21]. These subgraphs show where existing definitions violate the desired properties. Numerical label adjacent to each vertex denote its ID. The example in (B) contains a cyclic subgraph. It does not satisfy the definitions of ultrabubble and superbubble. The example in (C) is classified as a bubble by some definitions despite containing vertices that do not lie on any walk from the source to the sink. The example in (D) contains two overlapping structures that are classified as bubbles by some definitions.

Our main contributions are as follows:

- We introduce *panbubbles*, a subgraph abstraction that extends existing bubble definitions while retaining desirable structural properties in the presence of cycles (Figure 1A). Building on Mwaniki *et al*. [23], we also define *hairpins* to represent inversions flanked by inverted repeats.
- We prove that panbubbles and hairpins satisfy strict nesting, providing theoretical guarantees for their use in representing genomic variation.
- We develop an *O*(| *V* |^2^ (|*V* |+| *E*|))-time algorithm to identify all panbubbles and hairpins. We also propose a faster *O*(| *V* | (|*V* |+| *E*|))-time heuristic algorithm that produces the same output on all tested graphs. We implemented both algorithms in our tool `Billi`.
- On the largest publicly available human pangenome graph [21], which contains over 100 million vertices, `Billi` runs in under 75 minutes using 90 GB of memory, whereas VG requires approximately 20 hours and 500 GB of memory to compute snarls.
- We systematically compare panbubbles with existing definitions across 14 publicly available pangenome graphs, highlighting where the definitions differ. Using biologically relevant examples, we demonstrate that panbubbles and hairpins capture complex genomic structures, including inversions, that are not adequately represented by existing definitions.

## 2 Notations and problem statement

### Biedged graph

*Bidirected graphs* [10] are widely used in genomics because they offer a convenient way to represent adjacency relationships between DNA sequences, while allowing traversal of a sequence in both forward and reverse orientations. In this study, we adopt the *biedged graph* representation of bidirected graphs, following the approach of previous works [19,27]. A biedged graph *G*_*b*_ = (*V*_*b*_, *E*_*b*_, *f*_*b*_) is an undirected, colored multigraph subject to the following three constraints.

- Function *f*_*b*_ : *E*_*b*_ → {`gray`, `black`} assigns either gray or black color to each edge.
- Every vertex of the graph is incident with exactly one black edge.
- Between any two vertices (which may be identical), there can be at most one gray edge.

We call two vertices *u, v* ∈ *V*_*b*_ (*u* ≠ *v*) *adjacent* if they are connected by at least one edge of any color. With the above definition, adjacent vertices *u* and *v* can have either (i) one gray edge, (ii) one black edge, or (iii) one black and one gray edge between them (Figures 2 and 3A). For every vertex *v* ∈ *V*_*b*_, we use 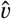 to denote the neighbor vertex of *v* connected using the unique black edge incident to *v*. We denote this black edge as *e*_*V*_. Clearly, 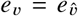 and 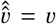 for all *v* ∈ *V*_*b*_. In genome sequence analysis, biedged graphs (or, equivalently, bidirected graphs) are used to represent genomic sequences and their adjacencies. Each black edge is annotated with a DNA sequence, while each gray edge represents an adjacency between two sequences. For a walk that traverses a black edge *e*_*V*_, the order in which vertices *v* and 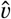 are visited determines the orientation (forward or reverse complement) in which the sequence label of *e*_*V*_ is read. In this work, we omit these sequence labels and focus solely on the underlying graph topology.

**Fig. 2:**
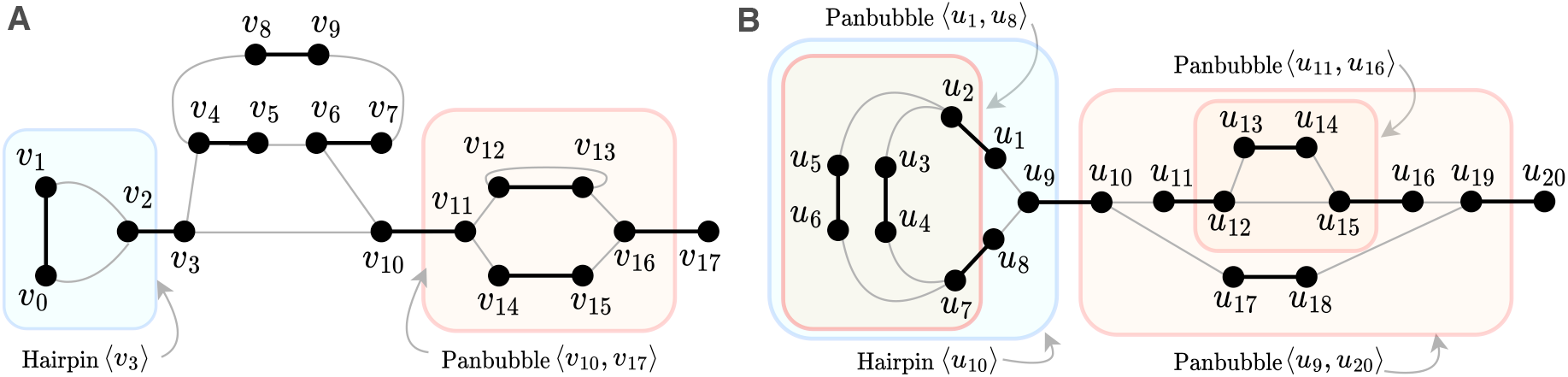
(A) Example of a hairpin and a panbubble is shown on the left. Hairpin ⟨*v*_3_⟩ is highlighted in a blue box. Panbubble ⟨*v*_10_, *v*_17_⟩ is highlighted in a red box. The blue box encloses all vertices in the set *Y*(*v*_3_) \ {*v*_3_}. The red box encloses all vertices in the set *B* (*v*_10_, *v*_17_). Note that the vertex pair (*v*_2_, *v*_11_) does not qualify as a panbubble entrance because vertices *v*_4_, *v*_5_, …, *v*_9_ violate the traversability condition. (B) The right figure illustrates the nesting relationships between panbubbles and hairpins. Panbubble ⟨*u*_11_, *u*_16_⟩ is nested within panbubble ⟨*u*_9_, *u*_20_⟩. Panbubble ⟨*u*_1_, *u*_8_⟩ is nested within hairpin ⟨*u*_10_⟩.

**Fig. 3:**
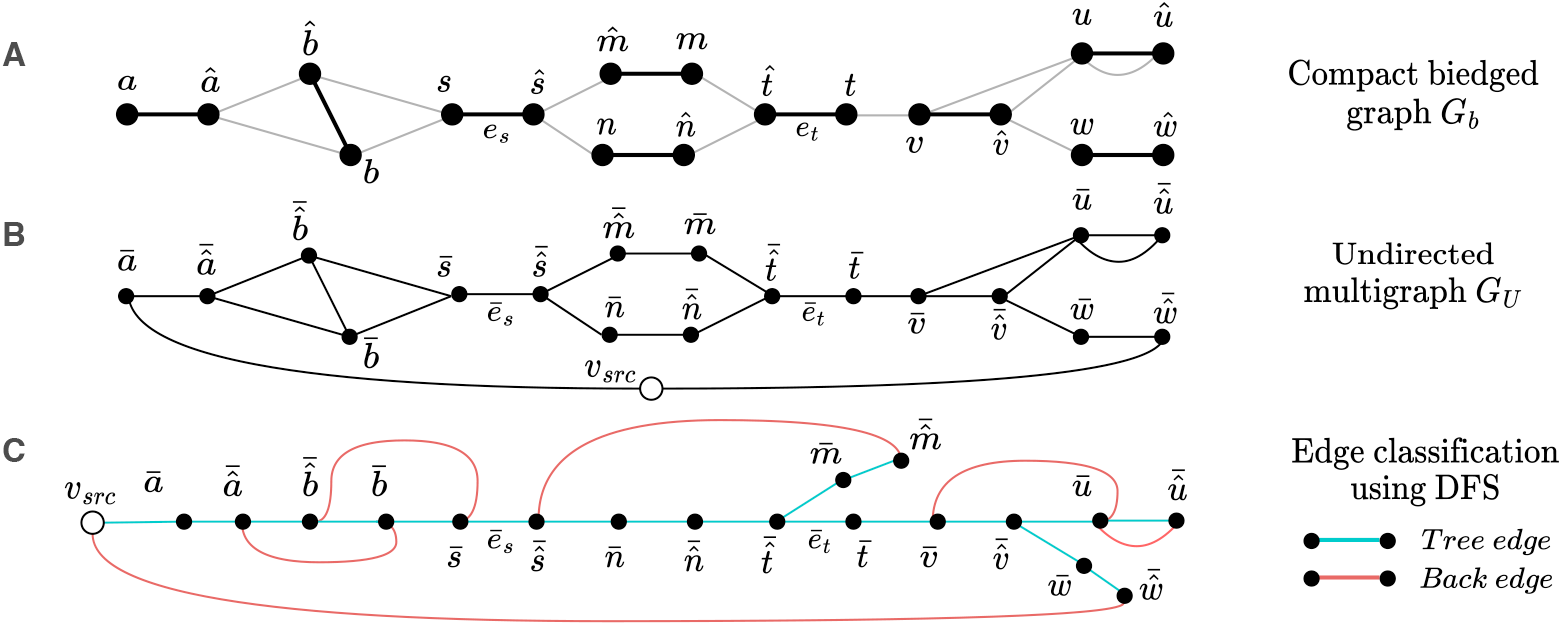
Illustration of the auxiliary graph construction. (A) Compact biedged graph *G*_*b*_ containing two panbubbles ⟨*a, ŝ*⟩ and ⟨*s, t*⟩. (B) The corresponding undirected multigraph *G*_*U*_. (C) Partitioning of the edges of *G*_*U*_ into tree edges and back edges using depth-first traversal.

The notion of a walk in a biedged graph differs from the standard definition of walks in undirected graphs. A sequence of vertices (*v*_0_, *v*_1_, …, *v*_*n*_) with *n* ≥ 1 is a *walk* in a biedged graph if (a) for all 1 ≤ *i* ≤ *n*, there exists an edge between *v*_*i*−1_ and *v*_*i*_, (b) the walk starts and ends with black edges, that is, 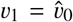 and 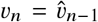, and (c) the entire walk alternates between black and gray edges. This definition reflects the requirement that we must finish reading the sequence associated with one black edge before starting the next. For example, vertex sequence (*v*_2_, *v*_3_, *v*_4_, *v*_5_, *v*_6_, *v*_7_) in Figure 2A forms a valid walk. On the other hand, the vertex sequence (*v*_2_, *v*_3_, *v*_4_, *v*_8_, *v*_9_) is not a walk because it contains two consecutive gray edges. Note that if vertex *v* appears in a walk, then its paired vertex 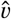 must also appear in that walk. Vertices and edges may be repeated in a walk. Moreover, walks are reversible: (*v*_*n*_, *v*_*n*−1_, …, *v*_1_, *v*_0_) is a walk whenever (*v*_0_, *v*_1_, …, *v*_*n*−1_, *v*_*n*_) is a walk.

Considering any *u, v* ∈ *V*_*b*_, we say that *v* is *reachable* from *u* if *v* appears in some walk starting from *u*. For example, in Figure 2A, vertices *v*_6_ and *v*_7_ are reachable from *v*_2_ because there exists a walk (*v*_2_, *v*_3_, *v*_4_, *v*_5_, *v*_6_, *v*_7_) starting from *v*_2_ containing *v*_6_ and *v*_7_. Observe that *v*_2_ is not reachable from *v*_6_ because no walk exists starting from *v*_6_ containing *v*_2_. By definition, the set of vertices reachable from *u* always includes *u* and *û* because (*u, û*) is a valid walk. We say that a walk (*v*_0_, *v*_1_, …, *v*_*n*−1_, *v*_*n*_) passes through *v*_*i*_ for all 1 ≤ *i* ≤ *n* − 1. Given any three vertices *u, v, w* ∈ *V*_*b*_, *v* is said to be reachable from *u* without passing through *w* if *v* appears in some walk starting from *u* and that walk does not pass through *w*. We call a vertex not incident with a gray edge a *tip*.

Two vertex-disjoint subgraphs of a biedged graph are said to be *disconnected* in the usual sense, that is, if there exists neither black edge nor gray edge that has one endpoint in the first subgraph and the other endpoint in the second subgraph. Unless a biedged graph has two or more disconnected subgraphs, it is said to be *connected*.

A walk (*v*_0_, *v*_1_, …, *v*_*n*−1_, *v*_*n*_) with *n* ≥ 3 is called a *closed walk* if *v*_*n*−1_ = *v*_0_ and *v*_*n*_ = *v*_1_. Similarly, a walk (*v*_0_, *v*_1_, …, *v*_*n*−1_, *v*_*n*_) with *n* 3 is an *inverted closed walk* if *v*_*n*−1_ = *v*_1_ and *v*_*n*_ = *v*_0_. For example, in Figure 2A, the vertex sequence (*v*_4_, *v*_5_, *v*_6_, *v*_7_, *v*_9_, *v*_8_, *v*_4_, *v*_5_) is a closed walk, whereas the vertex sequence (*v*_3_, *v*_2_, *v*_1_, *v*_0_, *v*_2_, *v*_3_) is an inverted closed walk.

### Compact biedged graph

Compacting long non-branching paths into single vertices is a common simplification in graph-based sequence analysis [5]. In the context of a biedged graph *G*_*b*_ = (*V*_*b*_, *E*_*b*_, *f*_*b*_), we define a *compaction operation* as follows. Consider two vertices *u, v* ∈ *V*_*b*_ with *u* ≠ *v* and 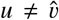. *u* and *v* are said to be *compactable* if (i) they are connected to each other by a gray edge and (ii) the number of gray edges incident to each of *u* and *v* is one. The compaction operation on a compactable vertex pair (*u, v*) consists of adding a black edge between *û* and 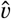, and then removing *u* and *v* together with all edges incident to either vertex. The resulting biedged graph has two fewer edges and two fewer vertices. A compact biedged graph is obtained by applying compaction operations as many times as possible in any order on the given biedged graph.

#### Definition 1 (Hairpin).

*Given a compact biedged graph G*_*b*_ = (*V*_*b*_, *E*_*b*_, *f*_*b*_), *let the set of vertices reachable from vertex s* ∈ *V*_*b*_ *without passing through s be denoted by Y* (*s*). *A hairpin is said to exist at vertex s if (i) every vertex v* ∈ *Y* (*s*) *appears in an inverted closed walk starting from s, (ii) removal of e*_*s*_, *i*.*e*., *the black edge connecting s and ŝ, separates the subgraph induced by Y* (*s*)\ {*s*} *from the rest of the graph, and (iii) no vertex in Y* (*s*) *other than s satisfies these conditions. The separable subgraph induced by Y* (*s*)\ {*s*} *is called a hairpin*.

For any vertex *s* that satisfies the above conditions, we refer to *s* and *e*_*s*_ as the *entrance vertex* and the *entrance edge* of the hairpin, respectively. This hairpin is denoted by ⟨*s*⟩.

#### Definition 2 (Panbubble).

*A panbubble exists between two vertices s and t (s ≠ t and* 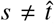*) in a compact biedged graph G*_*b*_ = (*V*_*b*_, *E*_*b*_, *f*_*b*_) *if all the following conditions hold:*

- matching: *Let U* (*s, t*) *denote the set of vertices reachable from s without passing through s or t. Analogously, define*
- *U* (*t, s*) *as the set of vertices reachable from t without passing through t or s. The matching condition requires that U* (*s, t*) = *U* (*t, s*).
- separable: *Define B* (*s, t*) *as U* (*s, t*)\{*s, t*}. *If black edges e*_*s*_ *and e*_*t*_ *are removed, the subgraph induced by B* (*s, t*) *is disconnected from the rest of the graph*.
- traversability: *Every vertex v* ∈ *U*(*s, t*) *appears on at least one walk from s to t*.
- no-hairpin: *No vertex in* 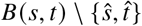 *is an entrance vertex of a hairpin*.
- minimality: *No vertex in U*(*s, t*) *other than t forms a pair* with *s and satisfies the above criteria. Similarly, no vertex in U*(*s, t*) *other than s forms a pair with t and satisfies the above criteria*.

The separable subgraph induced by *B* (*s, t*) is called a *panbubble*. For any set of two vertices {*s, t*} that satisfies these criteria, we denote the panbubble as ⟨*s, t*⟩ (or equivalently ⟨*t, s*⟩). In a panbubble ⟨*s, t*⟩, *s* and *t* are called *entrance vertices*. The edges *e*_*s*_ and *e*_*t*_ are called *entrance edges*. Note that the entrance vertices *s, t* and the entrance edges *e*_*s*_, *e*_*t*_ do not belong to the panbubble itself, i.e., they are not part of the subgraph induced by *B* (*s, t*). In Figure 2, we give examples of panbubbles and hairpins. Additional examples of valid and invalid subgraphs are provided in Supplementary Figure S2. These examples help clarify the role of the individual conditions in our definitions.

The panbubble definition is closely related to *bibubbles* [19]. Similar to a bibubble, a panbubble is not restricted to being acyclic. Moreover, both definitions include the matching, traversability, and separability conditions. The primary distinction between the two definitions is that we prohibit the occurrence of hairpins within panbubbles. We will show later in Theorems 3, 4, and 5 that this modification is essential to guarantee a strict nesting property among overlapping structures. In the next sections, we will address the following problem:

### Problem statement

Given a compact biedged graph *G*_*b*_ = (*V*_*b*_, *E*_*b*_, *f*_*b*_), (i) list the entrance vertex pairs of every panbubble in *G*_*b*_, and (ii) list the entrance vertices of every hairpin in *G*_*b*_.

## 3 Properties of panbubbles and hairpins

In this section, we present the key properties of panbubbles and hairpins, including their possible nesting relationships. These properties also inform the design of our detection algorithms for panbubbles and hairpins. For brevity, proofs of all claims in this paper are provided in the Supplementary Document. The following claim establishes upper bounds on the number of panbubbles and hairpins in a pangenome graph.

### Lemma 1.

*In a compact biedged graph G*_*b*_ = (*V*_*b*_, *E*_*b*_, *f*_*b*_), *the number of distinct panbubbles can be at most* |*V*_*b*_ |/2. *The number of distinct hairpins can be at most* |*V*_*b*_ |.

A valid walk in a biedged graph cannot traverse two gray edges consecutively, which limits the applicability of standard graph-theoretic techniques. To address this limitation, we construct an auxiliary undirected multigraph derived from *G*_*b*_ in which the edge colors are ignored, and the walk constraints are relaxed. The auxiliary graph will help us develop an efficient algorithm for panbubble and hairpin identification.

### Construction of an auxiliary undirected multigraph

We define our auxiliary undirected multigraph *G*_*U*_ = (*V*_*U*_, *E*_*U*)_ as follows. We assume that *G*_*b*_ is connected (if not, each connected component is handled independently). First, we copy all the edges and the vertices of *G*_*b*_ into *G*_*U*_. Note that tips in *G*_*b*_ correspond to vertices that have degree one in *G*_*U*_. We assume that *G*_*b*_ contains at least one tip^4^, and hence, *G*_*U*_ has at least one degree-one vertex. Next, we introduce one more vertex *v*_*src*_ in *G*_*U*_, and we connect *v*_*src*_ to all degree-one vertices (Fi<u>g</u>ures 3A, 3B). For every vertex *v* of *G*_*b*_, we denote the corresponding vertex in *G*_*U*_ by 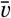. Vertex 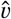 in *G*_*b*_ corresponds to 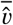 in *G*_*U*_. The edge of *G*_*U*_ corresponding to a black edge *e*_*V*_ in *G*_*b*_ is denoted by ē_*V*_. Finally, for every vertex *v V*∈_*b*_, we order the adjacency list of 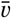 in *G*_*U*_ so that appears first. This ordering ensures that each edge in *G*_*U*_ derived from 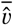 a black edge in *G*_*b*_ is explored as a tree edge, rather than a back edge, in any depth-first traversal of *G*_*U*_. The proof is straightforward and omitted for brevity.

Unlike biedged graphs, a walk in *G*_*U*_ is defined in the usual sense. An alternating sequence of vertices and edges, (*v*_0_, *e*_0_, *v*_1_, *e*_1_, *v*_2_,…, *e*_*n* −1_, *v*_*n*)_ with *n* ≥ 1 is a walk in *G*_*U*_ if edge *e*_*i*_ connects the vertices *v*_*i*_ and *v*_*i* +1_ for all 0 ≤ *i* ≤ *n* −1. A walk *v*_0_, *e*_0_, *v*_1_, *e*_1_, *v*_2_, …, *e*_*n* −1_, *v*_*n*_ passes through vertices *v*_1_, *v*_2_, …, *v*_*n*− 1_. A vertex is reachable from *v ∈V*_*U*_ if it appears in some walk starting from *v*. Path and cycle in *G*_*U*_ are also defined in the usual sense. A path is a walk whose vertices are distinct. A cycle in *G*_*U*_ is a closed path that starts and ends at the same vertex. We do not consider a self-loop as a cycle.

A depth-first traversal of *G*_*U*_, initiated at *v*_*src*_, partitions the edge set *E*_*U*_ into two classes: tree edges and back edges (Figure 3C). Let 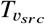 denote the spanning tree produced by this traversal, i.e., the subgraph of *G*_*U*_ that includes all vertices together with the tree edges. The tree 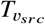 introduces a natural partial order *≺* on the vertices of *G*_*U*_. For *v*_1_, *v*_2_ *∈ V*_*U*_, *v*_1_ *≺ v*_2_ if and only if (i) *v*_1_ ≠ *v*_2_, (ii) *v*_1_ and *v*_2_ belong to a root-to-leaf path in 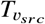, and (iii) *v*_1_ appears before *v*_2_ in that path. An analogous order is defined on the tree edges. For *e*_1_, *≺ e*_2_ *∈ E*_*U*_, we write *e*_1_ *e*_2_ if and only if (i)*e*_1_ ≠ *e*_2_, (ii) *e*_1_ and *e*_2_ belong to a root-to-leaf path in 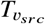, and (iii) *e*_1_ is traversed before *e*_2_ in that path. A vertex *w* in *V*_*U*_ is called an ancestor of a tree edge *e* = {*u, v*} with *u ≺ v* if either *w* = *u* or *w ≺ u*. Similarly, *w* is a descendant of edge *e* if either *w* = *v* or *v w*. Following Johnson et al. [16], we now introduce the notions of *bracket set* and *cycle equivalence*.

#### Definition 3 (Bracket, Bracket Set).

*A bracket of a tree edge e in G*_*U*_ *is a back edge connecting a descendant of e to an ancestor of e. The bracket set of a tree edge e is the set of all of its brackets*.

#### Definition 4 (Cycle-equivalent, Cycle-equivalence classes).

*Edges e*_1_ *and e*_2_ *are* cycle-equivalent *in G*_*U*_ *if every cycle in G*_*U*_ *contains either both edges or neither edge. The* cycle-equivalence class *of an edge e in G*_*U*_ *is the set of all edges that are cycle-equivalent to e*.

We now state several properties of panbubbles. For every panbubble ⟨*s, t*⟩ in *G*_*b*_, the vertices in *G*_*U*_ derived from *s, ŝ, t*, and 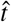 lie on a single root-to-leaf path of tree *T*_*vsrc*_ in a specific order (Lemma 2). Using the separable condition of ⟨*s, t*⟩, we show that edges *ē*_*s*_ and *ē*_*t*_ are cycle-equivalent (Lemma 3). The separable and traversability conditions of a panbubble ⟨*s, t*⟩ in *G*_*b*_ further imply that every interior black edge has one endpoint with a walk to *s* and the other with a walk to *t* (Lemma 4). The no-hairpin property of a panbubble implies that no interior black edge of ⟨*s, t*⟩ can correspond to a bracketless edge in *G*_*U*_, unless it lies on a common root-to-leaf path with *ē*_*s*_ and *ē*_*t*_ (Lemma 5).

#### Lemma 2.

*For every panbubble ⟨s, t* ⟩ *in G*_*b*_, *either* 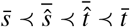 *or* 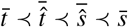 *in G*_*U*_.

#### Lemma 3.

*For every panbubble* ⟨*s, t* ⟩ *in G*_*b*_, *edges ē*_*s*_ *and ē*_*t*_ *are cycle equivalent in G*_*U*_.

#### Lemma 4.

*Suppose ⟨s, t*⟩ *is a panbubble in G*_*b*_. *Consider walks in G*_*b*_ *that end at s or t but avoid passing through s or t. Then, for every vertex u ∈U* (*s, t*), *there exist such walks from u to s and from* û*to t, or there exist such walks from* û *to s and from u to t*.

#### Lemma 5.

*Let* ⟨*s, t*⟩ *be a panbubble in G*_*b*_. *For every vertex* 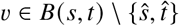, *if edge ē*_*V*_ *has an empty bracket set in G*_*U*_, *then either ē*_*s*_ *≺ ē*_*V*_ *≺ ē*_*t*_ *or ē*_*t*_ *≺ ē*_*V*_ *≺ ē*_*s*_.

Next we present Theorem 1 which states that the above conditions are both necessary and sufficient for a subgraph of *G*_*b*_ to represent a valid panbubble. We omit the minimality constraint in the claim because it will be enforced in our algorithm separately by processing candidate vertex pairs in a carefully chosen order.

#### Theorem 1.

*A vertex pair s, t ∈ V*_*b*_ *satisfies all the panbubble conditions (excluding minimality) if and only if all the following conditions hold:*

1. *Either* 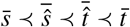 *or* 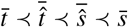 *in G*_*U*_.
2. *Edges ē*_*s*_ *and ē*_*t*_ *are cycle equivalent in G*_*U*_.
3. *Consider walks in G*_*b*_ *that end at s or t but avoid passing through s or t. Then, for every vertex u ∈ U* (*s, t*) ⋃ *U*(*t, s*), *there exist such walks from u to s and from* û *to t, or there exist such walks from û to s and from u to t*.
4. *For every vertex* 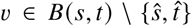, *if edge ē*_*V*_ *has an empty bracket set in G*_*U*_, *then either ē*_*s*_ *≺ ē*_*V*_ *≺ ē*_*t*_ *or ē*_*t*_ *≺ ē*_*V*_ *≺ ē*_*s*_.

Similarly, we state necessary and sufficient conditions for a subgraph of *NL* to represent a valid hairpin

#### Theorem 2.

*A vertex s is an entrance vertex of a hairpin in G*_*b*_ *if and only if all the following hold:*

1. *The bracket set of edge ē*_*s*_ *is empty in G*_*U*_.
2. *There does not exist any vertex v*∈*V*_*b*_ \ {*ŝ*} *such that* 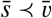 *in G*_*U*_ *and the bracket set of edge ē*_*V*_ *is empty*.
3. *Consider walks in G*_*b*_ *that end at s and avoid passing through s. For all vertices u*∈*Y* (*s*), *there exist such walks from both u and û*.

#### Nested relationships between panbubbles and hairpins

Pangenome graphs inherently support nested variation, as illustrated in Figure 2B. We characterize the conditions under which the subgraphs of *G*_*b*_ representing (i) two distinct hairpins, (ii) two distinct panbubbles, and (iii) a hairpin and a panbubble, may share a vertex. We first prove that if two panbubbles share a vertex, one must be fully contained within the other (Theorem 3). A panbubble and a hairpin can share a vertex only if the panbubble is completely contained in the hairpin (Theorem 4). Finally, two distinct hairpins can never share a vertex (Theorem 5).

##### Theorem 3.

*Let* ⟨*x, y*⟩ *and* ⟨*m, n*⟩ *be two different panbubbles such that B*(*x, y*)∩ *B*(*m, n*) ≠ ϕ, *then either B*(*x, y*) *is a strict subset of B*(*m, n*) *or B*(*m, n*) *is a strict subset of B*(*x, y*).

##### Theorem 4.

*Let* ⟨*x, y*⟩ *and* ⟨*z*⟩ *be a panbubble and a hairpin respectively in G*_*b*_ *such that B*(*x, y*)∩ (*Y* (*z*)\ {*z*}) ≠ ϕ, *then B*(*x, y*) *is a strict subset of Y* (*z*)\ {*z*}.

##### Theorem 5.

*Let* (*m*) *and* (*n*) *be two distinct hairpins in G*_*b*_. *Then, the two hairpins do not share any vertex, i*.*e*., (*Y* (*m*)\ {*m*}) ∩ (*Y* (*n*)\ {*n*}) = ϕ.

## 4 Algorithms to detect panbubbles and hairpins

### 4.1 Verifying a candidate panbubble entrance vertex pair

#### Notations and assumptions

Consider a candidate vertex pair (*s, t*) in *G*_*b*_. We assume that preprocessing has already confirmed that (*s, t*) satisfies the first and the second condition in Theorem 1. Details of this preprocessing will be discussed later in Section 4.3. We also assume that the following arrays have been precomputed:

- Boolean array *A*_*bridge*_ of size |*V*_*b*_ |such that *A*_*bridge*_ [*v*] = 1 if and only if the bracket set of edge *ē*_*V*_ is empty.
- Recall that we obtained spanning tree 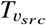 via a depth-first traversal of *G*_*U*_ (Section 3). Assume that arrays *A*_*d*_ and *A*_*f*_ record the discovery times and the finishing times of all vertices in *G*_*U*_, respectively. These arrays allow efficient checks of the ancestor-descendant relationships among vertices and edges in 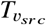[7].

#### Algorithm

We now describe how to verify the third and fourth conditions of Theorem 1. While evaluating a vertex pair (*s, t*) in *G*_*b*_, we use two boolean arrays *W*_*s*_ and *W*_*t*_, each of size |*V*_*b*_|. Array *W*_*s*_ will be used to mark those vertices that have walks originating from them to vertex *s* without passing through *s* or *t*. Similarly, array *W*_*t*_ will be used to mark those vertices that have walks originating from them to vertex *t* without passing through *s* or *t*.

We initialize arrays *W*_*s*_ and *W*_*t*_ with zeroes. First, we do a depth-first traversal starting from vertex *s* in *G*_*b*_. During this traversal, we visit all vertices that are reachable from *s* without passing through *s* or *t*. To respect the valid-walk restrictions in *G*_*b*_, we slightly modify the standard depth-first traversal procedure. Consider a single iteration of the traversal. Suppose the current vertex is *u* ∈ *V*_*b*_. We perform the following operations: (i) mark *u* as visited, (ii) update *W*_*s*_ [*û*] to 1, and then (iii) if *u* ≠ *ŝ* and 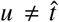, we recursively visit every unvisited neighbor of *û*. After the traversal completes, *W*_*s*_ [*v*] = 1 iff there exists at least one walk from *s* to *v* (or equivalently *v* to *s*) that does not pass through *s* or *t*, for all *v* ∈ *V*_*b*_. We repeat the same traversal starting from vertex *t* to compute array *W*_*t*_. We then declare the candidate pair (*s, t*) to satisfy all the conditions for a panbubble, except minimality, if both of the following conditions hold:

a. For every vertex 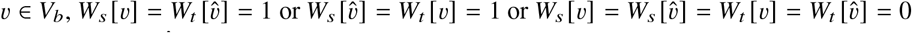.
b. For every vertex 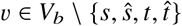 satisfying (i) *A*_*bridge*_ [*v*] = 1 and (ii) *W*_*s*_ [*v*] = 1 or *W*_*t*_ [*v*]= 1, the three edges *ē*_*s*_, *ē*_*V*_, *ē*_*t*_ lie on a common root-to-leaf path in 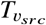.

### 4.2 Verifying a candidate hairpin entrance vertex

Consider a candidate entrance vertex *s* in *G*_*b*_. We will reuse the precomputed arrays *A*_*bridge*_, *A*_*d*_, and *A*_*f*_ defined in the previous section. To verify the three conditions of Theorem 2, we allocate a Boolean array *W* of size |*V*_*b*_|, initialized to zero. Array *W* will be used to mark those vertices that have walks originating from them to *s* without passing through *s*. To compute this array, we follow a depth-first traversal procedure customized for biedged graphs starting from vertex *s*. During this traversal, we visit all vertices that are reachable from *s* without passing through *s*. Consider a single iteration of the traversal. Suppose the current vertex is *u* ∈*V*_*b*_. We perform the following operations: (i) mark *u* as visited, (ii) update *W* [*û*] to 1, and then (iii) if *u* ≠ *ŝ*, we recursively visit every unvisited neighbor of *û*. After the traversal completes, *W* [*v*] = 1 iff there exists at least one walk from *s* to *v* (or equivalently *v* to *s*) that does not pass through *s*, for all *v* ∈*V*_*b*_. We declare vertex *s* as the entrance vertex of a hairpin if all the following conditions are satisfied:

a. *A*_*bridge*_ [*s*] = 1.
b. There is no vertex *v* ∈ *V*_*b*_ \ {*ŝ*} such that *A*_*bridge*_ [*v*] = 1 and 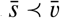 in *G*_*U*_.
c. For every vertex *v* ∈ *V*_*b*_, either 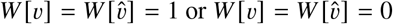.

### 4.3 Complete algorithms

We first construct the undirected multigraph *G*_*U*_ from the compact biedged graph *G*_*b*_ (Section 3). We next perform a standard depth-first traversal of *G*_*U*_ starting from *v*_*src*_ to obtain the spanning tree 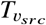 along with arrays *A*_*d*_ and *A*_*f*_. To compute array *A*_*bridge*_, we apply Tarjan’s linear-time bridge-finding algorithm [37]. Tarjan’s algorithm naturally extends to multigraphs by ensuring that parallel edges are handled explicitly, i.e., an edge is reported as a bridge only if no parallel edge provides an alternative connection between its endpoints.

#### Detection of panbubbles (exact algorithm)

We partition edges of *G*_*U*_ into cycle-equivalent classes by using the linear-time algorithm of Johnson *et al*. [16]. Let *C*_1_, *C*_2_, …, *C*_*k*_ denote the resulting classes, which we process one at a time. Consider a class *C*_*p*_, where 1 ≤ *p* ≤ *k*. Let *e*_1_, *e*_2_, …, *e*_*n*_ be the distinct black edges of *G*_*b*_ whose corresponding edges in *G*_*U*_ belong to *C*_*p*_. We enumerate these edges in level order of the DFS tree 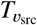. Accordingly, *ē*_*i*_ ≺ *ē*_*j*_ implies *i* < *j*. For each pair (*i, j*) with 1 ≤ *i* < *j* ≤ *n*, we check if *e*_*i*_ ≺ *e*_*j*_ holds. If so, we determine whether the vertex pair (*u, v*) satisfies the necessary conditions described in Section 4.1, where *e*_*i*_ = {*u, û*} with *u* ≺ *û* and 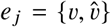 with 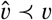. We examine these pairs using nested for loops: the outer loop iterates over *i* = 1, …, *n* − 1, and the inner loop iterates over *j* = *i* + 1, …, *n*. To enforce minimality, we terminate the inner loop as soon as a panbubble is detected.

#### Detection of panbubbles (heuristic algorithm)

We follow the same procedure as the exact algorithm, except that in the inner loop we restrict consideration to the first edge *e*_*k*_ such that *e*_*i*_ ≺*e*_*k*_, i.e., the edge with the smallest index *k* satisfying this condition. The remaining edges are not considered inside the inner loop. This design choice is based on our empirical finding that panbubbles rarely include internal edges that are cycle-equivalent to their entrance edges in *G*_*U*_. An example where this assumption would be violated is provided in Supplementary Figure S3.

#### Detection of hairpins (exact algorithm)

We iterate over all vertices *v* ∈*V*_*b*_ and check whether vertex *v* satisfies the necessary conditions discussed in Section 4.2. This hairpin-detection procedure can be paired with either the exact or heuristic panbubble-detection algorithm. The following theorem states the runtimes of our algorithms.

##### Theorem 6.

*Given a compact biedged graph G*_*b*_ = (*V*_*b*_, *E*_*b*_, *f*_*b*_), *the entrance vertices of all panbubbles and hairpins can be enumerated exactly in* O(|*V*_*b*_ |^2^ (|*V*_*b*_ |+ |*E*_*b*_ |)) *time. In contrast, the heuristic algorithm performs the same task in* O(|*V*_*b*_ |^2^(|*V*_*b*_ |+ |*E*_*b*_ |)) *time*.

## 5 Results

We implemented both the exact and heuristic algorithms in our software Billi, with the heuristic algorithm used by default. As demonstrated later, the heuristic algorithm significantly accelerates the computation without compromising accuracy. All experiments were done on a dual-socket Intel Xeon 6248R server with 2×24 cores and 750 GB RAM. **Benchmark graphs**. We used 14 publicly available pangenome graphs in GFA format, spanning a range of sizes and compositions. These graphs were constructed using six different graph construction tools: Minigraph [18], PGR-TK [6], Pangene [19], Bcalm2 [5], PGGB [11], and Minigraph-cactus [14]. These include graphs built from human, ape, and bacterial sequences. The size of each graph is reported in Supplementary Table S1, and the corresponding download links are listed in Supplementary Table S2. We preprocessed each graph to coalesce non-branching paths.

### Existing bubble detection methods

We computed panbubbles and hairpins using Billi. Superbubbles and ultrabubbles were computed using BubbleFinder [13]. We used VG to compute snarls. VG outputs a subset of snarls based on its filtering heuristics [27]. We computed bibubbles using Pangene [19]. The software versions and commands used are available in Supplementary Table S3. We did not include BubbleGun [8] in the comparison, as BubbleFinder is a more recent and faster method for superbubble identification [13]. We skipped povu [23] because we were unable to reliably interpret its output (github.com/pangenome/povu/issues/20).

### Bubble boundaries and nesting in a large inversion

A large fraction of bubbles in pangenome graphs are expected to correspond to common forms of sequence variation, such as substitutions, insertions, and deletions. These variants typically induce simple, acyclic bubbles that are well characterized and adequately detected with existing bubble definitions. A key motivation of this work is to relax the acyclicity constraint and enable the detection of complex structures that may correspond to biologically relevant events. For example, graph `MUC4` in our benchmark is derived from MUC4 gene sequences, which are known to contain variable-number tandem repeats with extensive allelic diversity [28]. The cyclic subgraph example shown earlier in Figure 1B in Section 1 was derived from `MUC4` graph.

Inversions are another class of complex variation that can be challenging to detect in pangenome graphs [31]. We investigated a known 46 kbp recurrent inversion in human chromosome Y, which is flanked by inverted repeats (Figure 4). Biological significance of recurrent inversions has been discussed previously in [29]. We observed a loop structure at the inversion event in graph `HPRCv1.1-chrY`. There also exist many SNPs in this region. The SNPs within the loop appear as nested variants in the graph. We then evaluated the output of the bubble detection methods in this region. The hairpin identified using Billi delineates the inverted region (Figure 4C). The panbubbles inside the hairpin correspond to nested variation within the inversion. In contrast, other bubble definitions do not capture this structure as effectively. VG reports incorrect variation boundaries and nesting relationships, whereas Pangene reports overlapping structures (Figure 4C). BubbleFinder does not detect the inversion because both the superbubble and ultrabubble definitions consider only acyclic subgraphs. In Billi, we avoid reporting overlapping panbubbles by enforcing the ‘no-hairpin’ constraint on panbubbles (Section 2) and reporting hairpins separately.

**Fig. 4:**
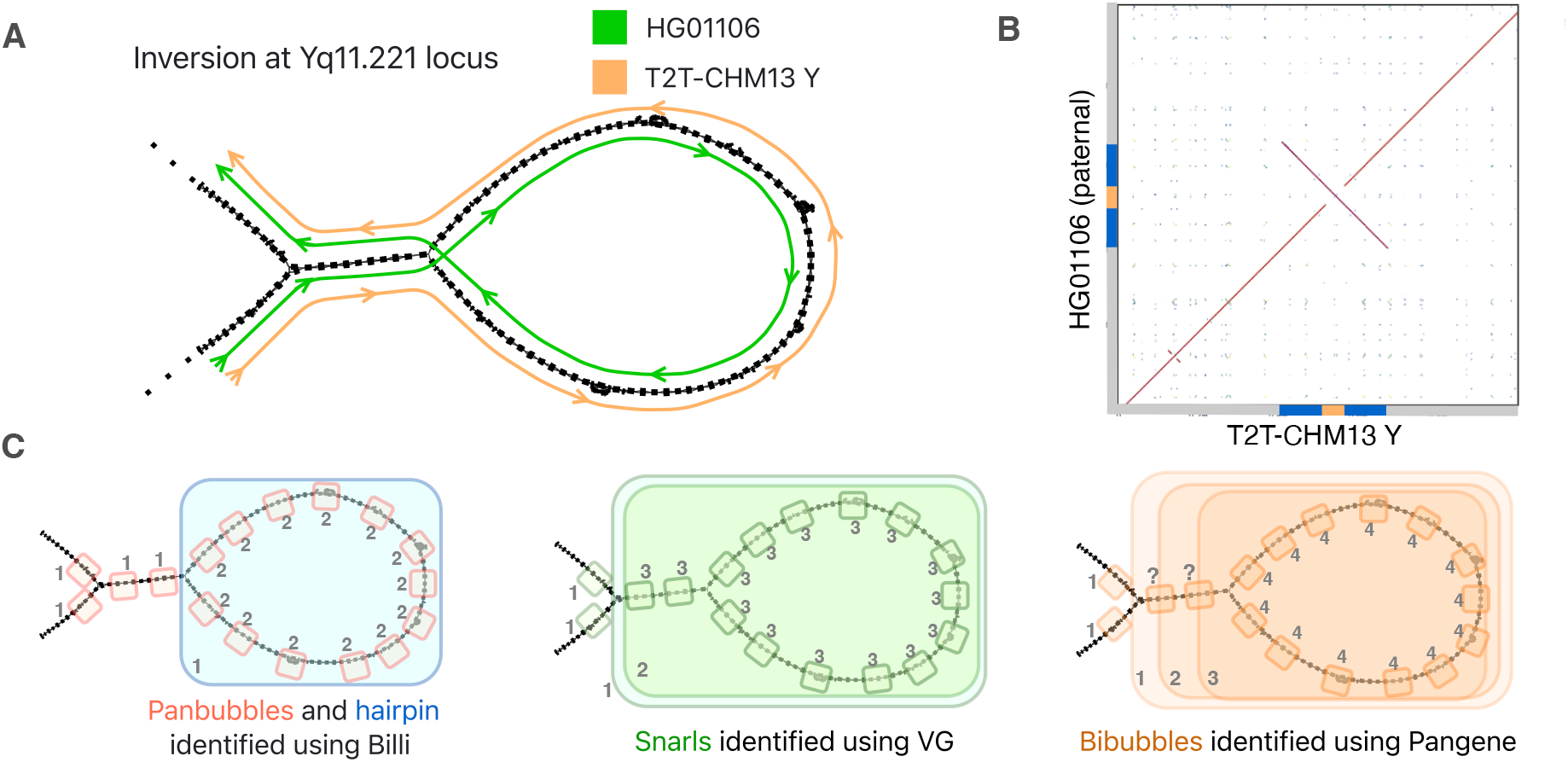
Illustrating the advantage of Billi using a complex inversion polymorphism in human chromosome Y. Chromosome Yq11.221 harbors a recurrent 46-kbp inversion flanked by repeats [29]. (A) Subgraph of `HPRCv1.1-chrY`, annotated with oppositely oriented path traversals for the HG01106 and T2T-CHM13 haplotypes. (B) Alignment dot plot between the corresponding assembly contigs, validating the inverted region. Flanking repeats and the inversion are highlighted in blue and gold, respectively. (C) Bubbles identified by Billi, VG, and Pangene, respectively. The numbers in small gray font indicate their nesting depth with ‘1’ indicating the top-level.

### Empirical comparison of bubble definitions

We next compare panbubbles with existing bubble definitions across all 14 graphs in our benchmark. The number of bubbles identified under the different definitions is reported in Table 1. For Billi, panbubble and hairpin counts are reported separately.

**Table 1:**
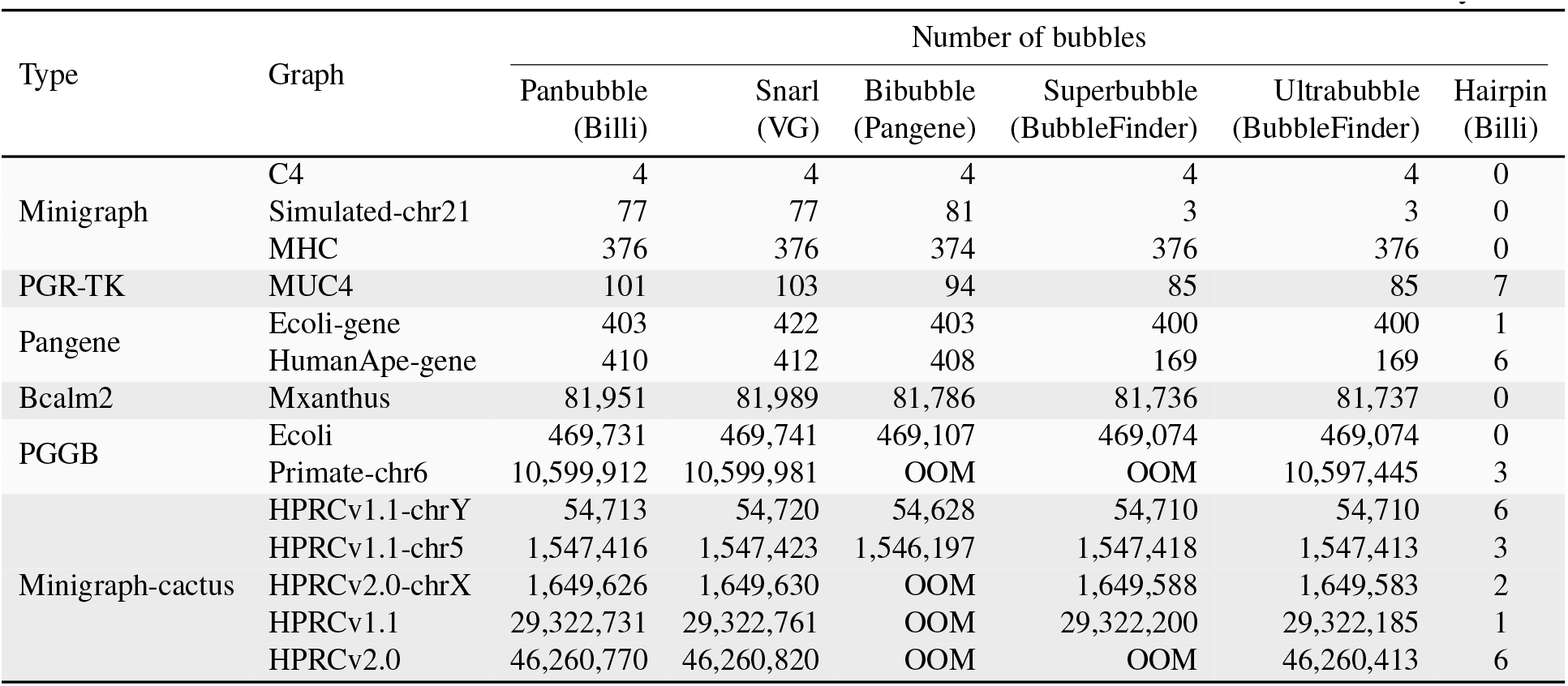
The number of bubbles detected under different bubble definitions. “OOM” indicates out-of-memory failure.

| Type | Graph | Number of bubbles |  |  |  |  |  |
| --- | --- | --- | --- | --- | --- | --- | --- |
|  |  | Panbubble<br>(Billi) | Snarl<br>(VG) | Bibubble<br>(Pangene) | Superbubble<br>(BubbleFinder) | Ultrabubble<br>(BubbleFinder) | Hairpin<br>(Billi) |
| Minigraph | C4 | 4 | 4 | 4 | 4 | 4 | 0 |
|  | Simulated-chr21 | 77 | 77 | 81 | 3 | 3 | 0 |
|  | MHC | 376 | 376 | 374 | 376 | 376 | 0 |
| PGR-TK | MUC4 | 101 | 103 | 94 | 85 | 85 | 7 |
| Pangene | Ecoli-gene | 403 | 422 | 403 | 400 | 400 | 1 |
|  | HumanApe-gene | 410 | 412 | 408 | 169 | 169 | 6 |
| Bcalm2 | Mxanthus | 81,951 | 81,989 | 81,786 | 81,736 | 81,737 | 0 |
| PGGB | Ecoli | 469,731 | 469,741 | 469,107 | 469,074 | 469,074 | 0 |
|  | Primate-chr6 | 10,599,912 | 10,599,981 | OOM | OOM | 10,597,445 | 3 |
| Minigraph-cactus | HPRCv1.1-chrY | 54,713 | 54,720 | 54,628 | 54,710 | 54,710 | 6 |
|  | HPRCv1.1-chr5 | 1,547,416 | 1,547,423 | 1,546,197 | 1,547,418 | 1,547,413 | 3 |
|  | HPRCv2.0-chrX | 1,649,626 | 1,649,630 | OOM | 1,649,588 | 1,649,583 | 2 |
|  | HPRCv1.1 | 29,322,731 | 29,322,761 | OOM | 29,322,200 | 29,322,185 | 1 |
|  | HPRCv2.0 | 46,260,770 | 46,260,820 | OOM | OOM | 46,260,413 | 6 |

Among the existing bubble definitions, recall that panbubbles, snarls, and bibubbles permit cyclic subgraphs, while superbubbles and ultrabubbles do not (Figure 1). This distinction is evident in the count of the bubbles across all graphs. The number of superbubbles and ultrabubbles is less than or equal to the counts reported by the other definitions. The difference is more pronounced in the `Simulated-chr21` and `HumanApe-gene` graphs. For example, graph `Simulated-chr21` contains many cyclic subgraphs associated with inversion events that are neither superbubble nor ultrabubble (Supplementary Figure S4A). The difference in bubble counts is less pronounced for Minigraph-cactus graphs, likely because Minigraph-cactus represents most repeat expansions as insertions rather than duplications [24]. Snarls are characterized solely by separability and minimality conditions [27]. VG performs additional filtering to conserve the nesting structure and to avoid partially overlapping snarls. Unlike panbubbles, the snarl definition permits the bubble subgraph to contain tips (Supplementary Figure S4B) and does not enforce the traversability condition (Figure 1C). This is a primary reason why VG often reports additional structures than Billi. We also encountered a few subgraphs for which we could not reason why VG did not report them as snarls (Supplementary Figure S4C).

Among the existing definitions, the bibubble definition used by Pangene [19] is the closest to panbubble. Bibubbles enforce traversability and do not allow tips. Pangene, however, employs a heuristic to reduce runtime that occasionally cause it to miss some bibubbles. Pangene examines only pairs of cycle-equivalent edges within a default distance of 100. Despite this heuristic, Pangene does not scale to the full HPRC graphs. Pangene and VG also identify a few bubbles containing hairpins that Billi does not report as panbubbles. As mentioned earlier, Billi reports hairpins separately.

Although bubble counts provide a useful high-level comparison, the counts do not convey whether different methods identify common or distinct structures. To assess this overlap, we also report the Jaccard similarity between the sets of bubble boundaries identified using each pair of definitions for three representative graphs in Supplementary Figure S5.

We also compared the outputs of Billi against the exact version of our algorithm (Section 4.3). The outputs of the two algorithms were identical for the first ten graphs. This result supports the rationale for our algorithmic optimization presented in Section 4.3, i.e., panbubbles tend to occur within the closest pair of cycle equivalent edges under the ≺ ordering. The exact algorithm did not scale to the remaining four graphs, including the full HPRC graphs, because it evaluates every pair of cycle equivalent edges. Overall, our experiments did not reveal any real instance in which the exact and heuristic algorithms produced different outputs. Nonetheless, developing a faster exact algorithm for panbubble detection that scales to large graphs remains an important direction for future work.

### Runtime, memory usage, and scalability to large pangenome graphs

We next evaluated the runtime and memory usage of all bubble detection methods (Table 2). Billi reports panbubbles and hairpins together in a single output file; consequently, its runtime includes the time required to compute both. Billi and Pangene were run sequentially because neither implementation supports multithreading. VG and BubbleFinder were run using 48 threads, corresponding to the number of cores available on our server.

**Table 2:**
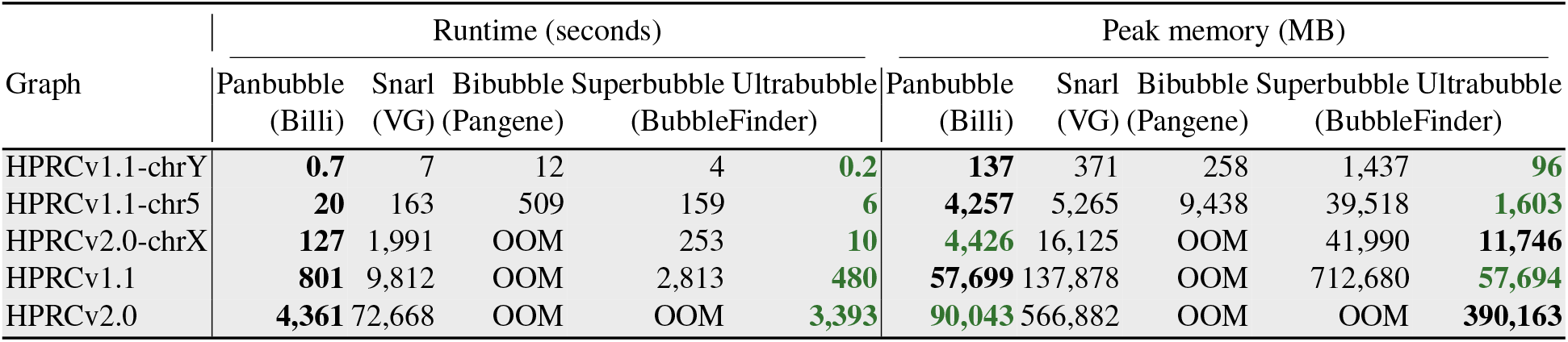
Runtime and memory usage of bubble detection methods. Results for smaller graphs are given in Supplementary Table S4. “OOM” indicates out-of-memory failure. **Green bold** = best, **black bold** = second best.

We could compute panbubbles, hairpins, ultrabubbles, and snarls for all 14 benchmark graphs. Bibubbles and superbubbles could not be computed on the largest graph `HPRCv2.0`. Among all bubble definitions, BubbleFinder had the lowest runtime for computing ultrabubbles, with Billi being a close second (e.g., 57 versus 73 minutes, respectively, on the largest graph `HPRCv2. 0`). For `HPRCv2. 0`, Billi required 90 GB of memory, less than the 390 GB required by `BubbleFinder` to compute ultrabubbles. Using comparable resources as BubbleFinder, Billi can identify panbubbles containing both cyclic or acyclic subgraphs. Compared with VG, Billi is significantly faster and memory efficient. Further optimization in Billi should be possible by processing chromosome-specific connected components in parallel. The heuristic algorithm used by Billi has a quadratic runtime complexity in the number of vertices. In contrast, BubbleFinder computes ultrabubbles and superbubbles using linear-time algorithms [13,32,33]. Nevertheless, Billi scales well to large pangenome graphs. Pangenome graphs tend to have small width in practice [22,30], which likely contributes to Billi’s practical scalability. Our heuristic algorithm performs graph traversals between the closest pairs of cycle-equivalent edges (Section 4.3); most traversals span only a small fraction of the graph in practice.

On the other hand, our exact algorithm did not finish computation in 24 hours for 4 out of 14 graphs. A performance comparison of the heuristic and exact algorithms is provided in Supplementary Table S5. The exact algorithm considers every pair of cycle-equivalent edges and therefore does not benefit from the same locality exploited by the heuristic approach. Lastly, for interested readers, we report the size distributions of panbubbles, and the counts of panbubbles at different nesting depths in Supplementary Tables S6 and S7, respectively.

## 6 Discussion

Pangenome-based variant detection promises to address the longstanding problem of reference bias. The draft human pangenome graph [20], for example, contains 119 Mbp of sequence not represented in the GRCh38 reference. As a step toward realizing this promise, we introduced panbubbles and hairpins, two subgraph abstractions for enumerating bubble-like structures in pangenome graphs. We proved that both satisfy strict nesting, along with other structural properties. Unlike walks in directed or undirected graphs, walks in biedged graphs must alternate between black and gray edges. As a result, the proofs required careful case-by-case analysis. We also discussed an exact algorithm and a scalable heuristic for enumerating panbubbles and hairpins in large pangenome graphs. Our experiments indicate that panbubbles and hairpins can capture complex genomic variation, including inversions with nested variants.

Several directions associated with improving the proposed algorithms remain open. First, it is unknown whether panbubbles and hairpins can be detected faster than *O* (|*V*|^2^ (|*V*| + |*E*|)) time in the worst case. Second, our current algorithms assume that the input graph contains at least one tip (Section 3). We leverage this property in our proofs because tips cannot occur within a panbubble or hairpin and therefore provide natural entry points for graph traversal. While this assumption holds for all the pangenome graphs in our benchmark, we encountered a graph from the Pangene study [19] containing a connected component with no tips. Our implementation currently aborts on such input. Developing algorithms that do not require this assumption is therefore an important direction.

Beyond algorithmic improvements, several challenges stand in the way of wider adoption of bubble-based approaches. First, the structures identified by any bubble-detection method depend on the underlying pangenome graph [31]. Thus, continued advances in genome assembly and pangenome graph construction are expected to improve the sensitivity and specificity of graph-based variant detection. Second, integrating pangenome bubbles into downstream analyses remains difficult. Standard formats such as VCF [9] describe variants relative to a single reference. Projecting bubbles onto a reference path loses their nesting and any non-reference structure [20]. Recent work has begun to address this limitation [24]. Third, benchmark resources from the Genome in a Bottle Consortium have been instrumental in developing and evaluating variant callers for linear reference genomes. Comparable well-characterized benchmarks for pangenome graphs would similarly accelerate progress in this field.

## Acknowledgments

We thank Shahbaz Khan, Paul Medvedev, and Heng Li for providing valuable feedback in this project. We also thank the Human Pangenome Reference Consortium (HPRC) for openly sharing their data. This research is supported in part by funding from DBT/Wellcome Trust India Alliance (IA/I/23/2/506979) and Prime Minister’s Research Fellowship (PMRF). We used computing resources provided by the National Energy Research Scientific Computing Center (NERSC), USA.

## Footnotes

4 In practice, pangenome graphs without tips are rare. Relaxing this assumption is left for future work.

