## Supplementary Document for "Billi: Accurate and Scalable Bubble Detection in Pangenome Graphs"

**Contents**


A. Supplementary figures ... 1
B. Supplementary tables ... 3
C. Properties of panbubbles and hairpins: full exposition ... 7
D. Algorithms to detect panbubbles and hairpins: full exposition ... 17


### A Supplementary figures

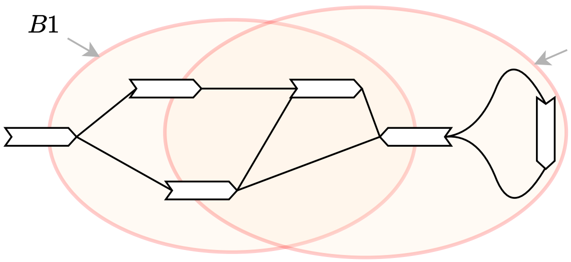


Fig. S1: Bubbles $B1$ and $B2$ in the above bidirected graph are valid snarls [27], bibubbles [19], and flubbles [23] by definition. Bubbles $B1$ and $B2$ overlap, i.e, the two subgraphs share vertices.

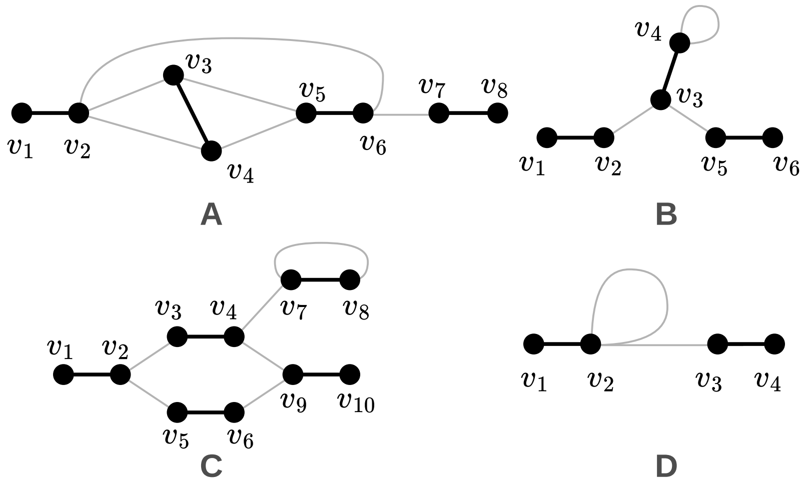


Fig. S2: **(A)** $\langle v_1, v_8 \rangle$ is a valid panbubble. However, $(v_1, v_6)$ is not a panbubble because it violates the separable condition. **(B)** $\langle v_3 \rangle$ is a valid hairpin; hence $(v_1, v_6)$ is not a panbubble because it violates the no-hairpin condition. **(C)** $(v_1, v_{10})$ is not a panbubble because $v_7$ and $v_8$ violate traversability. **(D)** $\langle v_1, v_4 \rangle$ is a valid panbubble.

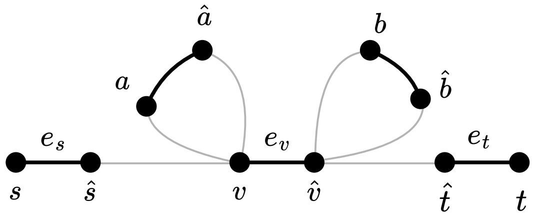


Fig. S3: An example where panbubble $\langle s, t\rangle$ contains an edge $e_v$ such that $\bar{e}_v, \bar{e}_s$, and $\bar{e}_t$ are cycle equivalent in $G_U$. Neither $(s, \hat{v})$ nor $(v, t)$ is a panbubble in this example, as each violates the traversability condition.

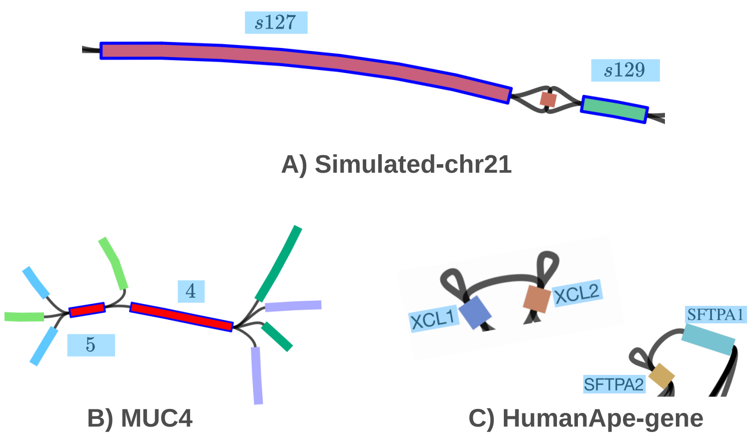


Fig. S4: **Bandage visualization of regions in pangenome graphs:** (A) Panbubble corresponding to an inversion variant in graph `Simulated-chr21`. It was also classified as a snarl and bibubble but not as a superbubble or ultrabubble. (B) A subgraph from graph `MUC4` containing a tip vertex between vertex 5 and vertex 4. It was classified as a snarl by VG. This was not classified as a panbubble. (C) Two subgraphs from graph `HumanApe-gene` that were not classified as snarls by VG. They were classified as panbubbles by Billi.

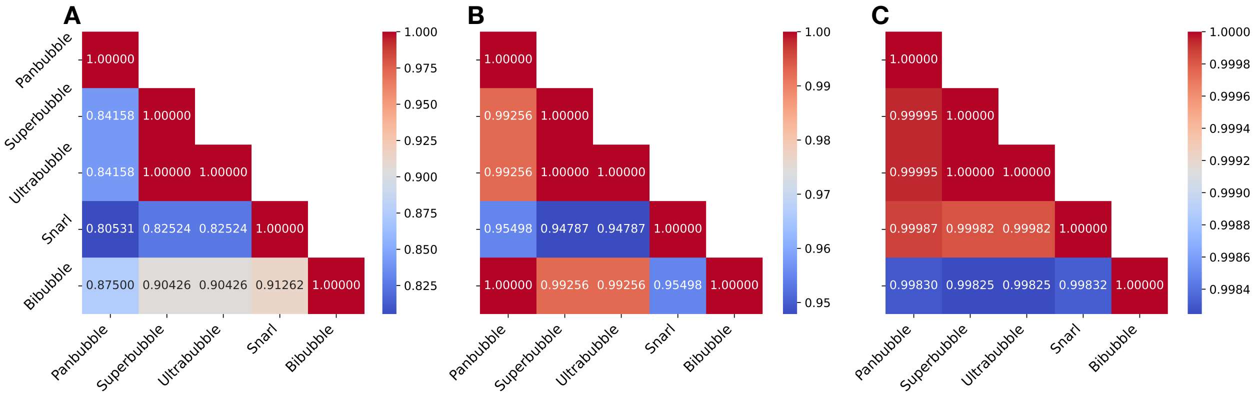


Fig. S5: Heatmaps showing the Jaccard similarity between the sets of bubbles identified by different bubble definitions for **(A)** `MUC4`, **(B)** `EColi-gene`, and **(C)** `HPRCv1.1-ChrY` graphs. These graphs were constructed using PGR-TK, Pangene, and Minigraph-cactus, respectively. The first column of each matrix shows the similarity between Panbubble and each of the other definitions. Similarity scores vary across the three graphs, likely due to differences in (a) the underlying sequence variation (e.g., bacterial genomes exhibit greater overall polymorphism and genomic diversity than human genomes) and (b) the graph-construction methodology (e.g., sequence alignment parameters and edge-selection heuristics). Each plot uses a different color scale.

### B Supplementary tables

Table S1: Publicly available pangenome graphs used in the experiments.

| Type | Graph | # Haplotypes | Before compaction | | After compaction | | # Components |
|---|---|---|---|---|---|---|---|
| | | | # Vertices | # Edges | # Vertices | # Edges | |
| Minigraph | C4 | 90 | 36 | 42 | 36 | 42 | 1 |
| | Simulated-chr21 | 10 | 316 | 466 | 316 | 466 | 1 |
| | MHC | 57 | 2,136 | 2,564 | 2,136 | 2,564 | 1 |
| PGR-TK | MUC4 | 97 | 6,374 | 9,251 | 4,170 | 5,563 | 1 |
| Pangene | Ecoli-gene | 50 | 26,018 | 51,875 | 12,344 | 18,769 | 22 |
| | HumanApe-gene | 110 | 38,082 | 63,925 | 4,162 | 7,664 | 1 |
| Bcalm2 | Mxanthus | 10 | 1,190,280 | 2,200,970 | 1,190,280 | 1,398,055 | 1 |
| PGGB | Ecoli | 50 | 3,107,334 | 3,673,827 | 3,107,030 | 3,673,523 | 1 |
| | Primate-chr6 | 14 | 68,531,256 | 81,333,257 | 68,531,198 | 81,333,199 | 9 |
| Minigraph-cactus | HPRCv1.1-chrY | 20 | 328,232 | 391,233 | 327,186 | 390,187 | 1 |
| | HPRCv1.1-chr5 | 90 | 9,946,742 | 11,894,826 | 9,943,370 | 11,891,454 | 1 |
| | HPRCv2.0-chrX | 377 | 10,434,952 | 12,437,642 | 10,268,330 | 12,271,020 | 1 |
| | HPRCv1.1 | 90 | 185,759,160 | 221,045,345 | 183,119,402 | 218,405,587 | 25 |
| | HPRCv2.0 | 464 | 296,566,820 | 354,315,094 | 291,459,944 | 349,208,218 | 25 |

**Description of graphs in Table S1.** The first three graphs `C4`, `Simulated-chr21`, and `MHC` were constructed using Minigraph [18]. Graph `C4` was constructed using 90 C4A/C4B gene sequences extracted from genome assemblies released by the Human Pangenome Research Consortium (HPRC) [20]. `Simulated-chr21` was constructed using ten human chromosome 21 sequences containing simulated substitution and inversion events [31]. Graph `MHC` was constructed using 57 Major Histocompatibility Complex (MHC) sequences. Minigraph excludes mutations smaller than 50 bp from its pangenome graphs, which is why its graphs will generally have fewer vertices and edges compared to other graph construction methods. The next graph `MUC4` was constructed from human MUC4 gene sequences using PGR-TK [6]. The next two graphs `Ecoli-gene` and `humanApe-gene` are *gene graphs* constructed using Pangene [19]. Graph `Ecoli-gene` was constructed using 50 *E. coli* genomes, whereas graph `humanApe-gene` was constructed using 100 human haplotypes and 10 great ape haplotypes [19]. Next, `Mxanthus` is a de Bruijn graph constructed using ten strains of *Myxococcus xanthus* [8]. Graphs `Ecoli` and `Primate-chr6` were constructed using PGGB [11]. The graphs `HPRCv1.1` and `HPRCv2.0` are the first and second human pangenome graph releases by HPRC, respectively, both using T2T CHM13 assembly [25] as the reference genome. Finally, graphs `HPRCv1.1-chrY` and `HPRCv1.1-chr5` are subgraphs of `HPRCv1.1` corresponding to chromosome Y and chromosome 5, respectively, and graph `HPRCv2.0-chrX` is the subgraph of `HPRCv2.0` corresponding to chromosome X.

Table S2: Graphs used in the benchmark and their corresponding download URLs. A subset of the graphs has been mirrored in a separate Zenodo repository: https://doi.org/10.5281/zenodo.21104719.

| Graph | URL |
|---|---|
| C4 | https://zenodo.org/records/6617246/files/C4-90.gfa.gz |
| Simulated-chr21 | https://zenodo.org/records/21170871/files/simulated_10hap.zip |
| MHC | https://zenodo.org/records/6617246/files/MHC-57.gfa.gz |
| MUC4 | https://giab-data.s3.amazonaws.com/PGR-TK-Files/CMRG_output_dir_v0.3.3.tar |
| Ecoli-gene | https://zenodo.org/records/14854301/files/Ecoli-v1.1-p0g30r5.gfa.gz |
| HumanApe-gene | https://zenodo.org/records/14854301/files/human100p10-v1.1-a1.gfa.gz |
| Mxanthus | https://github.com/fawaz-dabbaghieh/bubble_gun/blob/master/use_cases/m_xanthus_10_assemblies.unitigs.gfa.gz |
| Ecoli | https://zenodo.org/records/7937947/files/ecoli50.gfa.zst |
| Primate-chr6 | https://zenodo.org/records/7937947/files/primates14.chr6.fa.gz.667b9b6.c2fac19.ee137be.smooth.final.gfa.zst |
| HPRCv1.1-chrY | https://zenodo.org/records/15311999/files/chrY.zip |
| HPRCv1.1-chr5 | https://zenodo.org/records/15311999/files/chr5.zip |
| HPRCv2.0-chrX | https://s3-us-west-2.amazonaws.com/human-pangenomics/pangenomes/freeze/release2/minigraph-cactus/v2.0/hprc-v2.0-mc-chm13/hprc-v2.0-mc-chm13.chroms/chrX.vg |
| HPRCv1.1 | https://s3-us-west-2.amazonaws.com/human-pangenomics/pangenomes/freeze/freeze1/minigraph-cactus/hprc-v1.1-mc-chm13/hprc-v1.1-mc-chm13.gfa.gz |
| HPRCv2.0 | https://s3-us-west-2.amazonaws.com/human-pangenomics/pangenomes/freeze/release2/minigraph-cactus/v2.0/hprc-v2.0-mc-chm13/hprc-v2.0-mc-chm13.gfa.gz |

Table S3: Commands used to evaluate the bubble detection methods. For all tools, the input is a compact bidirected graph generated using Billi (`billi compact -s`). For Pangene, we additionally duplicated each edge in its reverse-complement orientation. In the `billi decompose` command, `-w` flag disables printing of alleles. For VG, `vg snarls` and `vg snarls -a` produced identical sets of compatible snarls. We used `vg snarls`, following the recommendation of the VG developers (github.com/vgteam/vg/issues/4957). Billi and Pangene do not support multithreading.

| Tool | Version | Threads | Command |
|---|---|---|---|
| Billi | v1.2 | 1 | billi decompose -i <INPUT_GFA_FILE> -w > <OUTPUT_FILE_PATH> |
| Billi (exact) | v1.2 | 1 | billi decompose -i <INPUT_GFA_FILE> -e -w > <OUTPUT_FILE_PATH> |
| VG | v1.75.1 | 48 | vg snarls -t 48 -n <INPUT_GFA_FILE> > <OUTPUT_FILE_PATH> |
| Pangene | commit:fc3366a | 1 | k8 pangene.js call -w <INPUT_GFA_FILE> > <OUTPUT_FILE_PATH> |
| BubbleFinder | v1.0.4 | 48 | BubbleFinder superbubbles -j 48 -g <INPUT_GFA_FILE> -o <OUTPUT_FILE_PATH><br>BubbleFinder ultrabubbles -j 48 -g <INPUT_GFA_FILE> -o <OUTPUT_FILE_PATH> |

Table S4: Runtime and peak memory required by bubble detection methods. "OOM" indicates termination due to insufficient memory. **Green bold** = best, **black bold** = second best.

| Graph | Runtime (seconds) | | | | | Peak memory (MB) | | | | |
|---|---|---|---|---|---|---|---|---|---|---|
| | Panbubble (Billi) | Snarl (VG) | Bibubble (Pangene) | Superbubble (BubbleFinder) | Ultrabubble | Panbubble (Billi) | Snarl (VG) | Bibubble (Pangene) | Superbubble (BubbleFinder) | Ultrabubble |
| C4 | **0.1** | **0.2** | **0.1** | **0.1** | **0.1** | **3** | 40 | **17** | 59 | 60 |
| Simulated-chr21 | **0.1** | 6 | **2** | **0.1** | **0.1** | **12** | 272 | **54** | 84 | 85 |
| MHC | **0.1** | **0.7** | **0.1** | **0.1** | **0.1** | **4** | 226 | **27** | 76 | 63 |
| MUC4 | **0.1** | **0.1** | **0.3** | 0.6 | **0.1** | **4** | 226 | **32** | 70 | 60 |
| Ecoli-gene | **0.1** | 0.3 | 0.4 | **0.2** | **0.1** | **8** | 139 | 67 | 70 | **61** |
| HumanApe-gene | **0.1** | 0.6 | **0.4** | **0.1** | **0.1** | **5** | 142 | 88 | **66** | 68 |
| Mxanthus | **5** | 16 | 103 | 20 | **0.8** | **449** | 699 | 1,063 | 4,471 | **202** |
| Ecoli | **6** | 32 | 125 | 51 | **2** | **1,206** | 1,897 | 2,963 | 13,071 | **493** |
| Primate-chr6 | **136** | 956 | OOM | OOM | **45** | **27,763** | 36,992 | OOM | OOM | **10,470** |
| HPRCv1.1-chrY | **0.7** | 7 | 12 | 4 | **0.2** | **137** | 371 | 258 | 1,437 | **96** |
| HPRCv1.1-chr5 | **20** | 163 | 509 | 159 | **6** | **4,257** | 5,265 | 9,438 | 39,518 | **1,603** |
| HPRCv2.0-chrX | **127** | 1,991 | OOM | 253 | **10** | **4,426** | 16,125 | OOM | 41,990 | **11,746** |
| HPRCv1.1 | **801** | 9,812 | OOM | 2,813 | **480** | **57,699** | 137,878 | OOM | 712,680 | **57,694** |
| HPRCv2.0 | **4,361** | 72,668 | OOM | OOM | **3,393** | **90,043** | 566,882 | OOM | OOM | **390,163** |

Table S5: Runtime and peak memory comparison of the heuristic and exact algorithms implemented in Billi. By default, the heuristic algorithm is used. "TLE" indicates time limit exceeded beyond 24 hours for those runs.

| Graph | Runtime (sec) | | Peak memory (MB) | |
|---|---|---|---|---|
| | Billi | Billi (exact) | Billi | Billi (exact) |
| C4 | 0.1 | 0.1 | 2.9 | 3 |
| Simulated-chr21 | 0.1 | 0.1 | 11.9 | 12.2 |
| MHC | 0.1 | 0.1 | 4.1 | 4.1 |
| MUC4 | 0.1 | 0.1 | 4.3 | 4.3 |
| Ecoli-gene | 0.1 | 0.1 | 7.9 | 8.0 |
| HumanApe-gene | 0.1 | 0.1 | 4.7 | 4.8 |
| Mxanthus | 5.1 | 5.4 | 449.2 | 450.1 |
| Ecoli | 6.2 | 6.4 | 1,206.0 | 1,204.8 |
| Primate-chr6 | 135.8 | 1,764.8 | 27,763.1 | 27,766.8 |
| HPRCv1.1-chrY | 0.7 | 67.4 | 136.6 | 137.5 |
| HPRCv1.1-chr5 | 20.2 | TLE | 4,256.6 | TLE |
| HPRCv2.0-chrX | 126.8 | TLE | 4,426.3 | TLE |
| HPRCv1.1 | 800.8 | TLE | 57,698.5 | TLE |
| HPRCv2.0 | 4,361.0 | TLE | 90,043.4 | TLE |

Table S6: Size statistics for all panbubbles and hairpins identified by Billi across the benchmark graphs. For each panbubble $\langle s, t \rangle$, we report its size as the number of vertices in its subgraph (i.e., $|B(s,t)|$). Likewise, the size of every hairpin $\langle m \rangle$ is given by the number of vertices in its corresponding subgraph excluding the entrance vertex (i.e., $|Y(m) \setminus \{m\}|$). Symbol "–" under the hairpin column denotes that no hairpins were found in that graph.

| | Cycle equivalence class size | | | Panbubble size | | | Hairpin size | | |
|---|---|---|---|---|---|---|---|---|---|
| Graph | Min | Median | Max | Min | Median | Max | Min | Median | Max |
| C4 | 3 | 3 | 3 | 6 | 6 | 28 | – | – | – |
| Simulated-chr21 | 78 | 78 | 78 | 4 | 4 | 6 | – | – | – |
| MHC | 2 | 2 | 207 | 4 | 6 | 496 | – | – | – |
| MUC4 | 2 | 2 | 39 | 2 | 10 | 194 | 1 | 1 | 1 |
| Ecoli-gene | 2 | 2 | 12 | 4 | 4 | 60 | 3 | 3 | 3 |
| HumanApe-gene | 2 | 3 | 22 | 2 | 4 | 58 | 1 | 1 | 3 |
| Mxanthus | 2 | 2 | 1,277 | 4 | 10 | 2,192 | – | – | – |
| Ecoli | 2 | 4 | 15,084 | 4 | 6 | 25,852 | – | – | – |
| Primate-chr6 | 2 | 3 | 1,671,453 | 4 | 6 | 935,054 | 1,485 | 7,991 | 68,263 |
| HPRCv1.1-chrY | 2 | 5 | 30,877 | 4 | 6 | 28,560 | 15 | 113 | 317 |
| HPRCv1.1-chr5 | 2 | 2 | 1,391,996 | 4 | 6 | 48,258 | 625 | 1,237 | 2,729 |
| HPRCv2.0-chrX | 2 | 2 | 1,390,824 | 4 | 6 | 163,132 | 3 | 3 | 11 |
| HPRCv1.1 | 2 | 3 | 1,986,080 | 4 | 6 | 1,436,456 | 747 | 747 | 747 |
| HPRCv2.0 | 2 | 2 | 2,815,823 | 4 | 6 | 2,293,044 | 3 | 11 | 2,259 |

Table S7: Count of panbubbles at different nesting levels. Level 1 is considered as the top level. Last column of the table indicates the deepest level until which panbubbles exist. Nested panbubbles may correspond to biological events or they may also arise as artifacts of pangenome graph construction and associated alignment heuristics [1].

| Graph | Count | | | | | | | Maximum depth |
|---|---|---|---|---|---|---|---|---|
| | Level 1 | Level 2 | Level 3 | Level 4 | Level 5 | Level 6 | Level > 6 | |
| C4 | 2 | 2 | 0 | 0 | 0 | 0 | 0 | 2 |
| Simulated-chr21 | 77 | 0 | 0 | 0 | 0 | 0 | 0 | 1 |
| MHC | 206 | 119 | 48 | 2 | 1 | 0 | 0 | 5 |
| MUC4 | 92 | 9 | 0 | 0 | 0 | 0 | 0 | 2 |
| Ecoli-gene | 383 | 20 | 0 | 0 | 0 | 0 | 0 | 2 |
| HumanApe-gene | 381 | 27 | 2 | 0 | 0 | 0 | 0 | 3 |
| Mxanthus | 55,606 | 24,753 | 1,554 | 38 | 0 | 0 | 0 | 4 |
| Ecoli | 346,384 | 101,207 | 19,709 | 2,227 | 202 | 2 | 0 | 6 |
| Primate-chr6 | 6,512,376 | 3,469,917 | 473,764 | 93,382 | 32,104 | 8,030 | 10,339 | 13 |
| HPRCv1.1-chrY | 30,875 | 15,563 | 7,054 | 996 | 212 | 13 | 0 | 6 |
| HPRCv1.1-chr5 | 1,412,357 | 96,987 | 28,865 | 7,536 | 1,339 | 218 | 114 | 8 |
| HPRCv2.0-chrX | 1,397,555 | 177,005 | 46,924 | 17,204 | 7,363 | 2,727 | 848 | 12 |
| HPRCv1.1 | 23,249,413 | 4,149,597 | 1,499,038 | 339,098 | 67,117 | 12,581 | 5,887 | 68 |
| HPRCv2.0 | 36,464,695 | 7,018,753 | 2,068,029 | 535,693 | 131,313 | 31,641 | 10,646 | 23 |

### C Properties of panbubbles and hairpins: full exposition

#### Proof of Lemma 1

Before proving Lemma 1, we first establish auxiliary results that will be useful in its proof.

**Lemma C.1.** *For every panbubble $\langle s,t\rangle$ in $G_b$, there exists a walk from $s$ to $t$ in which $s$ and $t$ appear only at the start and the end of the walk, respectively.*

*Proof.* Based on how we defined reachability in $G_b$ (Section 2), we know $t \in U(t,s)$. Also, $U(t,s) = U(s,t)$ due to the matching condition of $\langle s,t\rangle$. Therefore, $t \in U(s,t)$, which makes $t$ reachable from $s$ using some walk that does not pass through $s$ or $t$. This is only possible if there exists a walk from $s$ to $t$ in which $s$ and $t$ appear only at the start and the end of the walk, respectively.

**Lemma C.2.** *In a compact biedged graph $G_b = (V_b, E_b, f_b)$, a vertex can be the entrance vertex for at most one panbubble.*

*Proof.* We prove the claim by contradiction. Suppose there exist two distinct panbubbles $\langle s,t_1\rangle$, $\langle s,t_2\rangle$, where $s,t_1,t_2 \in V_b$ and $t_1 \neq t_2$.

First consider the case where $t_2 \in U(s,t_1)$. In this case, $t_2$ must also lie in $B(s,t_1)$. However, this contradicts the minimality condition of panbubble $\langle s,t_1\rangle$. Similarly, if $t_1 \in U(s,t_2)$, then $t_1$ lies in $B(s,t_2)$, contradicting the minimality condition of $\langle s,t_2\rangle$. Therefore, we must have $t_2 \notin U(s,t_1)$ and $t_1 \notin U(s,t_2)$.

Using Lemma C.1 on panbubble $\langle s,t_2\rangle$, there exists a walk $\omega$ from $s$ to $t_2$ in which $s$ and $t_2$ appear only at the start and the end of the walk, respectively. Since $t_2$ appears in $\omega$ and $t_2 \notin U(s,t_1)$, not all vertices in walk $\omega$ belong to $U(s,t_1)$. Consider the vertices of $\omega$ in the order starting from $s$. Let $v$ be the first vertex in the order that does not belong to $U(s,t_1)$. Since $v \notin U(s,t_1)$, walk $\omega$ must be of the form $(s,\dots,s,\dots,v,\dots,t_2)$ or $(s,\dots,t_1,\dots,v,\dots,t_2)$, i.e., the portion of walk $\omega$ between the start vertex $s$ and vertex $v$ must pass through $s$ or $t_1$. But it cannot pass through $s$ because $s$ appears only once in the entire walk. Therefore, walk $\omega$ must be of the form $(s,\dots,t_1,\dots,v,\dots,t_2)$. However, the appearance of $t_1$ in $\omega$ implies that $t_1$ is in $U(s,t_2)$. This violates the earlier conclusion that $t_1 \notin U(s,t_2)$.

**Lemma 1.** *In a compact biedged graph $G_b = (V_b, E_b, f_b)$, the number of distinct panbubbles can be at most $|V_b/2|$. The number of distinct hairpins can be at most $|V_b|$.*

*Proof.* A vertex cannot be the entrance vertex for two different panbubbles (Lemma C.2). In a compact biedged graph, there are $|V_b|$ vertices, but each panbubble is defined by two entrance vertices. Therefore, the number of panbubbles can be at most $|V_b/2|$.

Hairpins are defined by a single entrance vertex. Each vertex in $V_b$ can correspond to at most one hairpin. Therefore the number of distinct hairpins can be at most $|V_b|$.

#### Proof of Lemma 2

Recall the procedure of computing the auxiliary undirected multigraph $G_U$ and its depth-first spanning tree $T_{v_{src}}$ (Section 3). Every edge of $G_U$ must be either a tree edge or a back edge. The following corollary follows directly from the fundamental properties of depth-first traversal.

**Corollary C.1.** *For all $\{v_1,v_2\} \in E_U$ such that $v_1 \neq v_2$, either $v_1 \prec v_2$ or $v_2 \prec v_1$.*

Furthermore, every edge of $G_U$ derived from a black edge of $G_b$ must be a tree edge.

**Corollary C.2.** *For all $v \in V_b$, there exists a root-to-leaf path in $T_{v_{src}}$ that contains $\bar{v}$, $\bar{\bar{v}}$ and edge $\bar{e}_v = \{\bar{v},\bar{\bar{v}}\}$.*

For a panbubble $\langle s,t\rangle$ in $G_b$, let $\overline{U}(s,t)$ denote the set of vertices in $G_U$ corresponding to the set of vertices $U(s,t)$ in $G_b$. Formally, $\overline{U}(s,t) = \{\bar{v} | v \in U(s,t)\}$. Similarly, we define $\overline{B}(s,t) = \{\bar{v} | v \in B(s,t)\}$. Unlike the biedged graphs, reachability in $G_U$ is defined in the standard graph-theoretic sense (Section 3). The following property is therefore noteworthy.

**Lemma C.3.** *For each panbubble $\langle s,t\rangle$ in $G_b$, the set of vertices in $G_U$ reachable from $\bar{\bar{s}}$ without passing through $\bar{s}$ or $\bar{t}$ equals $\overline{U}(s,t)$.*

*Proof.* Let $X$ denote the set of vertices reachable from $\bar{\hat{s}}$ without passing through $\bar{s}$ or $\bar{t}$ in $G_U$. We need to show that $X = \overline{U}(s,t)$. We first argue that $\overline{U}(s,t) \subseteq X$. Recall that for every $v \in U(s,t)$, $v$ is reachable from $s$ in $G_b$ without passing through $s$ or $t$ (from the definition of $U(s,t)$). The corresponding walk in $G_U$ contains a subwalk from $\bar{\hat{s}}$ to $\bar{v}$ that does not pass through $\bar{s}$ or $\bar{t}$. Therefore, $\bar{v} \in X$ and $\overline{U}(s,t) \subseteq X$.

Next, we prove $X \subseteq \overline{U}(s,t)$ by contradiction. Suppose $X \not\subseteq \overline{U}(s,t)$. This implies the existence of a vertex in $G_U$ such that (i) this vertex is reachable from $\bar{\hat{s}}$ without passing through $\bar{s}$ or $\bar{t}$, and (ii) this vertex is not in $\overline{U}(s,t)$.

Under the above assumption, we can identify a walk $\omega$ having vertex sequence $(\bar{\hat{s}}, v_1, v_2, \ldots, v_k)$ in $G_U$ with $k \geq 1$ such that $\bar{s}$ and $\bar{t}$ do not appear in $\omega$, and $v_k$ is the only vertex in $\omega$ that does not belong to $\overline{U}(s,t)$ (if there are more vertices besides $v_k$ which do not belong to $\overline{U}(s,t)$, then one can trivially trim the walk to ensure that only the last vertex is not in $\overline{U}(s,t)$). It follows that vertices $v_1, v_2, \ldots, v_{k-1}$ are in $\overline{U}(s,t) \setminus \{\bar{s}, \bar{t}\} = \overline{B}(s,t)$. Further note that $v_k \neq v_{src}$, because if $v_k = v_{src}$, then we would have a tip vertex among $B(s,t)$, which contradicts the traversability condition of $\langle s,t \rangle$.

Consider the case when $v_k \neq v_{src}$ and $k > 1$. Since $v_{k-1} \in \overline{B}(s,t)$ and $v_k \neq v_{src}$, we must have vertices corresponding to $v_{k-1}$ and $v_k$ in $G_b$. We must also have an edge in $G_b$ connecting the two vertices. Since $v_{k-1} \in \overline{B}(s,t)$ and $v_k \notin \overline{U}(s,t)$, observe that the presence of an edge in $G_b$ between a vertex in $B(s,t)$ and a vertex in $V_b \setminus U(s,t)$ contradicts the separable condition of $\langle s,t \rangle$.

The other case, i.e., $v_k \neq v_{src}$ and $k = 1$, implies the presence of an edge in $G_b$ between $\hat{s}$ and a vertex in $V_b \setminus U(s,t)$, again contradicting the separable condition of $\langle s,t \rangle$.

The arguments in the above proof also imply the following.

**Corollary C.3.** *For each panbubble $\langle s,t \rangle$ in $G_b$, the set of vertices in $G_U$ reachable from $\bar{\hat{s}}$ without passing through $\bar{s}$ or $\bar{t}$ does not contain $v_{src}$.*

Next, we analyze the properties of edges $\bar{e}_s$ and $\bar{e}_t$ in $G_U$, which are derived from panbubble entrance edges $e_s$ and $e_t$ in $G_b$.

**Lemma C.4.** *For each panbubble $\langle s,t \rangle$ in $G_b$, either $\bar{e}_s \prec \bar{e}_t$ or $\bar{e}_t \prec \bar{e}_s$ in $G_U$.*

*Proof.* By contradiction. Suppose neither $\bar{e}_s \prec \bar{e}_t$ nor $\bar{e}_t \prec \bar{e}_s$. Then there does not exist any root-to-leaf path in $T_{v_{src}}$ that contains both $\bar{e}_s$ and $\bar{e}_t$. Suppose wlog that $\bar{s}$ was marked as visited before $\bar{t}$ during the depth-first traversal of $G_U$ from vertex $v_{src}$.

By Corollary C.1, we have either $\bar{s} \prec \bar{\hat{s}}$ or $\bar{\hat{s}} \prec \bar{s}$. We consider the two cases one by one.

Case 1 ($\bar{s} \prec \bar{\hat{s}}$): During the depth-first traversal, $\bar{\hat{s}}$ must have been visited immediately after $\bar{s}$. From Lemma C.1, there exists a walk $(s, \hat{s}, \ldots, t)$ in $G_b$ not passing through $s$ or $t$. Analogously, there exists a walk $(\bar{s}, \bar{\hat{s}}, \ldots, \bar{t})$ in $G_U$ not passing through $\bar{s}$ or $\bar{t}$. This means vertex $\bar{t}$ (and thus edge $\bar{e}_t$) must lie within the sub-tree of $\bar{\hat{s}}$ in $T_{v_{src}}$ This contradicts our initial assumption that there is no root-to-leaf path in $T_{v_{src}}$ that contains both $\bar{e}_s$ and $\bar{e}_t$.

Case 2 ($\bar{\hat{s}} \prec \bar{s}$): During the depth-first traversal, $\bar{s}$ must be visited immediately after $\bar{\hat{s}}$ . Also, since $\bar{s}$ is marked as visited before $\bar{t}$ during the traversal, $\bar{t}$ is certainly not an ancestor of $\bar{e}_s$. As a result, there exists a path from $\bar{\hat{s}}$ to $v_{src}$ in $G_U$ without passing through $\bar{s}$ or $\bar{t}$. This contradicts our previous claim in Corollary C.3.

**Lemma C.5.** *For each panbubble $\langle s,t \rangle$ in $G_b$, $\bar{e}_s \prec \bar{e}_t$ if and only if $\bar{s} \prec \bar{\hat{s}} \prec \bar{\hat{t}} \prec \bar{t}$.*

*Proof.* $[\Leftarrow]$ $\bar{s} \prec \bar{\hat{s}} \prec \bar{\hat{t}} \prec \bar{t}$ directly implies $\bar{e}_s \prec \bar{e}_t$.

$[\Rightarrow]$ If $\bar{e}_s \prec \bar{e}_t$, then clearly $\bar{s} \prec \bar{t}$, $\bar{s} \prec \bar{\hat{t}}$, $\bar{\hat{s}} \prec \bar{t}$, and $\bar{\hat{s}} \prec \bar{\hat{t}}$. Next we will argue that $\bar{s} \prec \bar{\hat{s}}$. Assume for contradiction that $\bar{\hat{s}} \prec \bar{s}$. In that case, there would exist a path from $\bar{\hat{s}}$ to $v_{src}$ without passing through $\bar{s}$ or $\bar{t}$, which contradicts our earlier claim in Corollary C.3.

Next we prove that $\bar{\hat{t}} \prec \bar{t}$ by contradiction. Suppose $\bar{t} \prec \bar{\hat{t}}$. Then $\bar{s} \prec \bar{\hat{s}} \prec \bar{t} \prec \bar{\hat{t}}$ and there exists a path $p = (\bar{s}, \bar{\hat{s}}, \ldots, \bar{t})$ in $G_U$ consisting entirely of tree edges. By construction, this path does not include $\bar{\hat{t}}$. Vertices in a path are distinct by definition. Using Lemma C.3, every vertex on path $p$ belongs to $\overline{U}(s,t)$. Similarly, every vertex on path $p$ except $\bar{s}$ and $\bar{t}$ belongs to $\overline{B}(s,t)$. Based on this path in $G_U$, if we consider the corresponding sequence of vertices in $G_b$, then observe that vertex $t$ must connect to some vertex in $B(s,t)$ other than $\hat{t}$. This contradicts the separable condition of panbubble $\langle s,t \rangle$.

Using a symmetric argument, we obtain the following:

**Corollary C.4.** *For each panbubble $\langle s,t \rangle$ in $G_b$, $\bar{e}_t \prec \bar{e}_s$ if and only if $\bar{t} \prec \bar{\hat{t}} \prec \bar{\hat{s}} \prec \bar{s}$.*

**Lemma 2.** *For every panbubble $\langle s,t\rangle$ in $G_b$, either $\bar{s} \prec \bar{\hat{s}} \prec \bar{\hat{t}} \prec \bar{t}$ or $\bar{t} \prec \bar{\hat{t}} \prec \bar{\hat{s}} \prec \bar{s}$ in $G_U$.*

*Proof.* For a panbubble $\langle s,t\rangle$ in $G_b$, we know either $\bar{e}_s \prec \bar{e}_t$ or $\bar{e}_t \prec \bar{e}_s$ in $G_U$ (Lemma C.4). From Lemma C.5, and Corollary C.4, we have that either $\bar{s} \prec \bar{\hat{s}} \prec \bar{\hat{t}} \prec \bar{t}$ or $\bar{t} \prec \bar{\hat{t}} \prec \bar{\hat{s}} \prec \bar{s}$ in $G_U$.

#### Proof of Lemma 3

**Lemma C.6.** *Tree edges $e_1$ and $e_2$ are cycle equivalent in $G_U$ if and only if they have the same set of brackets.*

*Proof.* This property holds for all undirected multigraphs; see Theorem 5 in [16].

**Lemma C.7.** *For each panbubble $\langle s,t\rangle$ in $G_b$, edges $\bar{e}_s$ and $\bar{e}_t$ have the same bracket set in $G_U$.*

*Proof.* By contradiction. Assume that edges $\bar{e}_s$ and $\bar{e}_t$ have different bracket sets. Using Lemma C.4, either $\bar{e}_s \prec \bar{e}_t$ or $\bar{e}_t \prec \bar{e}_s$. Suppose wlog that $\bar{e}_s \prec \bar{e}_t$. Accordingly, $\bar{s} \prec \bar{\hat{s}} \prec \bar{\hat{t}} \prec \bar{t}$ (Lemma C.5). Let $D_s$ and $D_t$ denote the set of all descendant vertices of edges $\bar{e}_s$ and $\bar{e}_t$ in tree $T_{v_{src}}$, respectively. Let $A_s$ and $A_t$ denote the set of all ancestor vertices of the edges $\bar{e}_s$ and $\bar{e}_t$, respectively. For $\bar{e}_s$ and $\bar{e}_t$ to have an unequal bracket set, an edge must exist in $G_U$ such that either (a) one endpoint of the edge is in $D_s \setminus D_t$ and the other endpoint is in $A_s$, or (b) one end-point of the edge is in $D_t$ and the other endpoint is in $A_t \setminus A_s$.

Suppose the edge satisfies condition (a). By Corollary C.3, the endpoint of this edge in $A_s$ cannot be $v_{src}$. Now, observe that using this edge, we have a path between $\bar{\hat{s}}$ and $\bar{s}$ in $G_U$ without using $\bar{e}_s$ or $\bar{e}_t$. Mapping this path back to the corresponding sequence of edges in $G_b$ contradicts the separable condition of $\langle s,t\rangle$. Similarly, if the edge satisfies Condition (b), then there exists a path between $\bar{\hat{s}}$ and $\bar{t}$ in $G_U$ without using $\bar{e}_s$ or $\bar{e}_t$. Considering the corresponding sequence of edges in $G_b$ again contradicts the separable condition of $\langle s,t\rangle$.

**Lemma 3.** *For every panbubble $\langle s,t\rangle$ in $G_b$, edges $\bar{e}_s$ and $\bar{e}_t$ are cycle-equivalent in $G_U$.*

*Proof.* Edges $\bar{e}_s$ and $\bar{e}_t$ have the same bracket set (Lemma C.7). Hence, these two edges are also cycle equivalent in $G_U$ (Lemma C.6).

#### Proof of Lemma 4

**Lemma C.8.** *Suppose $\langle s,t\rangle$ is a panbubble in biedged graph $G_b$. If $u \notin U(s,t)$ and $v \in U(s,t)$, then every walk from $u$ to $v$ includes $s$ or $t$.*

*Proof.* By contradiction. Assume that there exists a walk between $u$ and $v$ containing neither $s$ nor $t$. Note that $u \neq s$ and $u \neq t$ because $s,t \in U(s,t)$. If either $v = s$ or $v = t$ holds, then we are done. Therefore, we consider the case when $u \notin U(s,t)$ and $v \in B(s,t)$. The walk between $u$ and $v$ does not traverse the black edges $e_s$ or $e_t$ (because otherwise it would include $s$ or $t$). This further implies that removing the two entrance edges of the panbubble does not disconnect the subgraph induced by $B(s,t)$ from the subgraph induced by $V_b \setminus B(s,t)$. This contradicts the separable condition of $\langle s,t\rangle$.

The following claim holds directly from the above lemma.

**Corollary C.5.** *Suppose $\langle s,t\rangle$ is a panbubble in a biedged graph. If $u \notin B(s,t)$ and $v \in B(s,t)$, then every walk from $u$ to $v$ includes $s$ or $t$.*

**Lemma 4.** *Suppose $\langle s,t\rangle$ is a panbubble in $G_b$. Consider walks in $G_b$ that end at $s$ or $t$ but avoid passing through $s$ or $t$. Then, for every vertex $u \in U(s,t)$, there exist such walks from $u$ to $s$ and from $\hat{u}$ to $t$, or there exist such walks from $\hat{u}$ to $s$ and from $u$ to $t$.*

*Proof.* For convenience, let us call a walk *special* if it ends at $s$ or $t$ and does not pass through $s$ or $t$. Consider a vertex $u \in U(s,t)$. When $u \in \{s,\hat{s},t,\hat{t}\}$, the lemma holds trivially (Lemma C.1). Therefore, let us consider the case when $U(s,t) \setminus \{s,\hat{s},t,\hat{t}\} \neq \phi$ and $u \in U(s,t) \setminus \{s,\hat{s},t,\hat{t}\}$.

From the matching condition of $\langle s,t\rangle$, we know that $u$ is reachable from $s$ without passing through $s$ or $t$, and $u$ is reachable from $t$ without passing through $s$ or $t$. Accordingly, by the definition of reachability in $G_b$, there exist special walks from $u$ or $\hat{u}$ to $s$, and there exist special walks from $u$ or $\hat{u}$ to $t$. Accordingly, there are four possible combinations:

(1) There exist special walks from $u$ to $s$ and from $u$ to $t$,
(2) There exist special walks from $\hat{u}$ to $s$ and from $\hat{u}$ to $t$,
(3) There exist special walks from $u$ to $s$ and from $\hat{u}$ to $t$, and
(4) There exist special walks from $\hat{u}$ to $s$ and from $u$ to $t$,

where at least one of the above must be true. If (3) or (4) holds, then the lemma follows directly.

We will next prove that if (2) holds, then (3) or (4) must also hold. Suppose, for the sake of contradiction, that (2) holds, while neither (3) nor (4) holds. Then there exist special walks from $\hat{u}$ to $s$ and from $\hat{u}$ to $t$. However, there exists no special walk from $u$ to $s$ or $t$.

From the traversability condition of $\langle s, t\rangle$, there exists a walk $\omega$ from $s$ to $t$ in which vertices $u$ and $\hat{u}$ appear. One of the following two sub-cases must hold: Either (a) the first occurrence of $u$ immediately precedes the first occurrence of $\hat{u}$ in $\omega$, or (b) the first occurrence of $u$ immediately follows the first occurrence of $\hat{u}$ in $\omega$.

Suppose case (a) holds. Let us consider the sub-walk $\omega'$ of $\omega$ from the the first occurrence of $u$ to the last vertex of $\omega$, i.e., $t$. From our earlier conclusion, recall that there is no special walk from $u$ to $t$. Therefore, $\omega'$ must pass through vertices $s$ or $t$. Consider the first pass of this walk through either black edge $e_s$ or $e_t$. The corresponding ordered pair of vertices that appear in $\omega'$ must be one of $(s, \hat{s})$, $(\hat{s}, s)$, $(t, \hat{t})$ or $(\hat{t}, t)$. Among these pairs, observe that pairs $(\hat{s}, s)$ or $(\hat{t}, t)$ cannot occur because they would imply the existence of a special walk from $u$ to $s$ or $t$. On the other hand, if the pair is $(s, \hat{s})$, then this would lead to violation of the separable condition of $\langle s, t\rangle$. This is because the vertex immediately preceding $s$ in $\omega'$ cannot be $\hat{s}$, and it must be in $B(s, t)$ (Corollary C.5). The presence of a gray edge between a vertex in $B(s, t) \setminus \{\hat{s}\}$ and $s$ violates the separable condition of $\langle s, t\rangle$. An analogous argument can be used to show that pair $(t, \hat{t})$ cannot occur. This leads to a contradiction, which confirms that if (2) holds, then (3) or (4) must also hold.

Now suppose case (b) holds, where the first occurrence of $u$ immediately follows the first occurrence of $\hat{u}$ in $\omega$. Recall that walks in a biedged graph are symmetric, i.e., any walk from one vertex to another can be reversed to obtain a walk in the opposite direction. Let us consider the sub-walk $\omega'$ of $\omega$ from the the first occurrence of $u$ in $\omega$ to the first vertex of $\omega$, i.e., $s$. One can now repeat similar arguments as case (a) to arrive at a contradiction. Therefore, if (2) holds, then (3) or (4) must also hold.

The above proof can be easily extended to show that if (1) holds, then (3) or (4) must also hold.

**Proof of Lemma 5**

**Lemma C.9.** *Consider a panbubble $\langle s, t\rangle$ in $G_b$ such that $\bar{e}_s \prec \bar{e}_t$. For all $u \in U(s, t)$, there exists a root-to-leaf path containing $\bar{s}$ and $\bar{u}$ in $T_{v_{src}}$.*

*Proof.* Using Lemma C.5, we know that $\bar{s} \prec \bar{\hat{s}} \prec \bar{\hat{t}} \prec \bar{t}$. Let us assume for contradiction that there exists a vertex $u \in U(s, t)$ such that there is no root-to-leaf path containing $\bar{s}$ and $\bar{u}$. Since $u \in U(s, t)$, $u$ is reachable from $s$ without passing through $s$ or $t$ in $G_b$. Accordingly, in $G_U$, there exists a walk with vertex sequence $(\bar{s}, \bar{\hat{s}}, \ldots, \bar{u})$ that does not pass through $\bar{s}$ or $\bar{t}$. There also exists a path from $\bar{u}$ to $v_{src}$ that includes neither $\bar{s}$ nor $\bar{t}$. This means $v_{src}$ is reachable from $\bar{\hat{s}}$ without passing through $\bar{s}$ or $\bar{t}$, contradicting our claim in Corollary C.3.

**Lemma C.10.** *Consider a panbubble $\langle s, t\rangle$ in $G_b$ such that $\bar{e}_s \prec \bar{e}_t$. Let $D_s$ and $D_t$ denote the set of all descendant vertices of edges $\bar{e}_s$ and $\bar{e}_t$, respectively. For all $v \in V_b$, $\bar{v} \in D_s \setminus D_t$ if and only if $v \in B(s, t)$.*

*Proof.* [$\Rightarrow$] Using Lemma C.5, we have $\bar{s} \prec \bar{\hat{s}} \prec \bar{\hat{t}} \prec \bar{t}$. If $\bar{v} \in D_s \setminus D_t$, then there exists a path between $\bar{\hat{s}}$ and $\bar{v}$ without passing through $\bar{s}$ or $\bar{t}$. Using Lemma C.3, it follows that $\bar{v} \in \overline{U}(s, t)$. Therefore, $v \in U(s, t)$. Furthermore, $\bar{v}$ cannot be $\bar{s}$ or $\bar{t}$ because $\bar{s}, \bar{t} \notin D_s \setminus D_t$. As a result, $v \in B(s, t)$.

[$\Leftarrow$] Assume for contradiction that $\bar{v} \notin D_s \setminus D_t$. Since $v \in B(s, t)$, (i) by the definition of $B(s, t)$ in $G_b$, there exists a walk $\omega$ with vertex sequence $(\bar{s}, \bar{\hat{s}}, \ldots, \bar{v})$ in $G_U$ which does not pass through $\bar{s}$ or $\bar{t}$, and (ii) there exists a root-to-leaf path in $T_{v_{src}}$ containing $\bar{s}$ and $\bar{v}$ (Lemma C.9). Since $\bar{v} \notin D_s \setminus D_t$ and $\bar{v}$ is neither $\bar{s}$ nor $\bar{t}$, $\bar{v}$ must be either an ancestor of $\bar{s}$ or a descendant of $\bar{t}$. Since walk $\omega$ from $\bar{s}$ to $\bar{v}$ does not pass through $\bar{s}$ or $\bar{t}$, the bracket sets of $\bar{e}_s$ and $\bar{e}_t$ must be unequal. This contradicts our claim in Lemma C.7.

**Corollary C.6.** *Consider a panbubble $\langle s, t\rangle$ in $G_b$ such that $\bar{e}_s \prec \bar{e}_t$. For all $u \in B(s, t)$, $\bar{s} \prec \bar{u}$ in $G_U$.*

**Lemma 5.** *Let $\langle s, t\rangle$ be a panbubble in $G_b$. For every vertex $v \in B(s, t) \setminus \{\hat{s}, \hat{t}\}$, if edge $\bar{e}_v$ has an empty bracket set in $G_U$, then either $\bar{e}_s \prec \bar{e}_v \prec \bar{e}_t$ or $\bar{e}_t \prec \bar{e}_v \prec \bar{e}_s$.*

*Proof.* By contradiction. Suppose there exists a vertex $v \in B(s,t) \setminus \{\hat{s}, \hat{t}\}$ such that, in $G_U$: (a) $\bar{e}_v$ has an empty bracket set, and (b) neither $\bar{e}_s \prec \bar{e}_v \prec \bar{e}_t$ nor $\bar{e}_t \prec \bar{e}_v \prec \bar{e}_s$ holds. We will show that the existence of such a vertex $v$ implies the presence of a hairpin entrance vertex in $B(s,t) \setminus \{\hat{s}, \hat{t}\}$, thereby violating the no-hairpin condition of $\langle s, t \rangle$.

Suppose wlog that $\bar{v} \prec \bar{\hat{v}}$. From Lemma C.4, either $\bar{e}_s \prec \bar{e}_t$ or $\bar{e}_t \prec \bar{e}_s$. Assume wlog that $\bar{e}_s \prec \bar{e}_t$. Let $D_s, D_v$ and $D_t$ denote the set of all descendant vertices of edges $\bar{e}_s, \bar{e}_v$, and $\bar{e}_t$ respectively. Since $v \in B(s,t) \setminus \{\hat{s}, \hat{t}\}$, it follows that $\bar{e}_s \prec \bar{e}_v$ and $\bar{v} \in D_s \setminus D_t$ (using Corollary C.6 and Lemma C.10). Since $\bar{e}_s \prec \bar{e}_v \prec \bar{e}_t$ does not hold, edges $\bar{e}_v$ and $\bar{e}_t$ do not share a root-to-leaf path, which means $D_v \subseteq \overline{B}(s,t) \setminus \{\bar{\hat{s}}, \bar{\hat{t}}, \bar{v}\}$.

As the bracket set of $\bar{e}_v$ is empty, removing $\bar{e}_v$ would disconnect the vertices in $D_v$ from the rest of $G_U$. The rest of $G_U$ includes $\bar{s}$ and $\bar{t}$. The same property should hold in $G_b$, that is, the removal of $e_v$ would disconnect the subgraph induced by vertex set $\{u | u \in V_b, \bar{u} \in D_v\}$ from the rest of the graph (which includes $s$ and $t$). For every vertex $u$ in this vertex set, the traversability condition of $\langle s, t \rangle$ ensures the existence of a walk from $s$ to $t$ passing through $u$. Thus, the vertex set $\{u | u \in V_b, \bar{u} \in D_v\}$ equals $Y(v) \setminus \{v\}$ and every vertex in $Y(v)$ must appear in an inverted closed walk starting from $v$. As a result, there must exist a hairpin with entrance vertex $v$ or with some other entrance vertex within $Y(v)$. Since $Y(v) \subseteq (B(s,t) \setminus \{\hat{s}, \hat{t}\})$, the no-hairpin condition of $\langle s, t \rangle$ is violated.

### Proof of Theorem 1

For a hairpin $\langle s \rangle$ in $G_b$, let $\overline{Y}(s)$ denote the set of vertices in $G_U$ corresponding to the set $Y(s)$ in $G_b$. Formally, $\overline{Y}(s) = \{\bar{v} | v \in Y(s)\}$. We establish a few intermediate results for hairpins before proving Theorem 1.

**Lemma C.11.** *For each hairpin $\langle s \rangle$ in $G_b$, the set of vertices in $G_U$ reachable from $\bar{\hat{s}}$ without passing through $\bar{s}$ equals $\overline{Y}(s)$.*

*Proof.* We follow the same proof technique used for Lemma C.3. Let $X$ denote the set of vertices in $G_U$ that are reachable from $\bar{\hat{s}}$ without passing through $\bar{s}$. We need to show that $X = \overline{Y}(s)$. We first argue that $\overline{Y}(s) \subseteq X$. Recall that for every $v \in Y(s)$, $v$ is reachable from $s$ without passing through $s$ (from the definition of $Y(s)$). The corresponding walk in $G_U$ contains a subwalk from $\bar{\hat{s}}$ to $\bar{v}$ that does not pass through $\bar{s}$. Therefore, $\bar{v} \in X$.

Next, we prove $X \subseteq \overline{Y}(s)$ by contradiction. Suppose that $X \not\subseteq \overline{Y}(s)$. This implies the existence of a vertex in $G_U$ such that (i) this vertex is reachable from $\bar{\hat{s}}$ without passing through $\bar{s}$, and (ii) this vertex is not in $\overline{Y}(s)$. In that case, there must exist at least one walk $\omega$ having vertex sequence $(\bar{\hat{s}}, v_1, v_2, \ldots, v_k)$ in $G_U$ with $k \geq 1$ such that $\bar{s}$ does not appear in $\omega$, and $v_k$ is the only vertex in $\omega$ that does not belong to $\overline{Y}(s)$. It follows that vertices $v_1, v_2, \ldots, v_{k-1}$ are in $\overline{Y}(s) \setminus \{\bar{s}\}$. Further note that $v_k \neq v_{src}$, because if $v_k = v_{src}$, then we would have a tip vertex in $Y(s) \setminus \{s\}$. But such a tip vertex cannot appear in an inverted closed walk starting from $s$, contradicting that $\langle s \rangle$ is a hairpin.

Consider the case when $v_k \neq v_{src}$ and $k > 1$. Since $v_{k-1} \in \overline{Y}(s) \setminus \{\bar{s}\}$ and $v_k \neq v_{src}$, we must have vertices corresponding to $v_{k-1}$ and $v_k$ in $G_b$. We must also have an edge in $G_b$ between the two vertices. Since $v_{k-1} \in \overline{Y}(s) \setminus \{\bar{s}\}$ and $v_k \notin \overline{Y}(s)$, observe that the presence of an edge in $G_b$ between a vertex in $Y(s) \setminus \{s\}$ and a vertex in $V_b \setminus Y(s)$ contradicts the separable property of hairpin $\langle s \rangle$.

The other case, i.e., $v_k \neq v_{src}$ and $k = 1$, implies the presence of an edge in $G_b$ between $\hat{s}$ and a vertex in $V_b \setminus Y(s)$, again contradicting the separable property of hairpin $\langle s \rangle$.

The arguments in the above proof also imply the following.

**Corollary C.7.** *For each hairpin $\langle s \rangle$ in $G_b$, the set of vertices in $G_U$ reachable from $\bar{\hat{s}}$ without passing through $\bar{s}$ does not contain $v_{src}$.*

Corollary C.7 further implies the following.

**Corollary C.8.** *For each hairpin $\langle s \rangle$ in $G_b$, $\bar{s} \prec \bar{\hat{s}}$ in $G_U$.*

**Lemma C.12.** *For each hairpin $\langle s \rangle$ in $G_b$, the bracket set of edge $\bar{e}_s$ in $G_U$ is empty.*

*Proof.* By contradiction. Assume that the bracket set of edge $\bar{e}_s$ is non-empty. From Corollary C.8, we have $\bar{s} \prec \bar{\hat{s}}$ in $G_U$. Let $D_s$ denote the set of all descendant vertices of edge $\bar{e}_s$. Let $A_s$ denote the set of all ancestor vertices of edge $\bar{e}_s$. As the bracket set of edge $\bar{e}_s$ is non-empty, there must exist a back edge such that one endpoint of the edge is in $D_s$ and the other endpoint of the edge is in $A_s$. From Corollary C.7, the endpoint of this edge in $A_s$ cannot be $v_{src}$. Now, observe that using this edge, we have a path between $\bar{\hat{s}}$ and $\bar{s}$ in $G_U$ without using $\bar{e}_s$. Mapping this path back to the corresponding sequence of edges in $G_b$ contradicts the separable property of hairpin $\langle s \rangle$.

**Lemma C.13.** *Consider a hairpin* $\langle s \rangle$ *in* $G_b$*. For all* $u \in Y(s)$*, there exists a root-to-leaf path containing* $\bar{s}$ *and* $\bar{u}$ *in* $T_{v_{src}}$*.*

*Proof.* Assume for contradiction that there exists a vertex $u \in Y(s)$ such that there is no root-to-leaf path containing $\bar{s}$ and $\bar{u}$. Since $u$ belongs to $Y(s)$, $u$ is reachable from $s$ without passing through $s$ in $G_b$. Accordingly, in $G_U$, there exists a walk with vertex sequence $(\bar{s}, \bar{\hat{s}}, \ldots, \bar{u})$ that does not pass through $\bar{s}$. Under our assumption that $\bar{s}$ and $\bar{u}$ do not share a root-to-leaf path, there also exists a path from $\bar{u}$ to $v_{src}$ that does not include $\bar{s}$. This means $v_{src}$ is reachable from $\bar{\hat{s}}$ without passing through $\bar{s}$, contradicting Corollary C.7.

**Lemma C.14.** *Consider a hairpin* $\langle s \rangle$ *in* $G_b$*. Let* $D_s$ *denote the set of all descendant vertices of edge* $\bar{e}_s$*. For all* $v \in V_b$*,* $\bar{v} \in D_s$ *if and only if* $v \in Y(s) \setminus \{s\}$*.*

*Proof.* [$\Rightarrow$] From Corollary C.8, we know that $\bar{s} \prec \bar{\hat{s}}$ in $G_U$. If $\bar{v} \in D_s$, then there exists a path from $\bar{\hat{s}}$ to $\bar{v}$ without passing through $\bar{s}$. Using Lemma C.11, it follows that $\bar{v} \in \overline{Y}(s)$. Therefore, $v \in Y(s)$. Furthermore, $\bar{v}$ cannot be $\bar{s}$ because $\bar{s} \notin D_s$. As a result, $v \in Y(s) \setminus \{s\}$.

[$\Leftarrow$] Assume for contradiction that there exists a vertex $v \in Y(s) \setminus \{s\}$ such that $\bar{v} \notin D_s$. Since $v \in Y(s) \setminus \{s\}$, (i) there exists a walk $\omega$ with vertex sequence $(\bar{s}, \bar{\hat{s}}, \ldots, \bar{v})$ in $G_U$ which does not pass through $\bar{s}$, and (ii) there exists a root-to-leaf path in $T_{v_{src}}$ containing $\bar{s}$ and $\bar{v}$ (Lemma C.13). Since $\bar{v} \notin D_s$ and $\bar{v} \neq \bar{s}$, $\bar{v}$ must be an ancestor of $\bar{s}$. Since walk $\omega$ from $\bar{s}$ to its ancestor vertex $\bar{v}$ does not pass through $\bar{s}$, the bracket set of edge $\bar{e}_s$ must contain a back edge, contradicting Lemma C.12.

**Corollary C.9.** *Consider a hairpin* $\langle s \rangle$ *in* $G_b$*. For all* $u \in Y(s) \setminus \{s\}$*,* $\bar{s} \prec \bar{u}$ *in* $G_U$*.*

**Lemma C.15.** *In a hairpin* $\langle s \rangle$ *in* $G_b$*, there does not exist any vertex* $v \in Y(s) \setminus \{s, \hat{s}\}$ *such that the bracket set of edge* $\bar{e}_v$ *in* $G_U$ *is empty.*

*Proof.* By contradiction. Suppose there exists a vertex $v \in Y(s) \setminus \{s, \hat{s}\}$ such that the bracket set of $\bar{e}_v$ in $G_U$ is empty. Suppose wlog that $\bar{v} \prec \bar{\hat{v}}$. From Corollary C.8, we have $\bar{s} \prec \bar{\hat{s}}$. From Corollary C.9, we have $\bar{s} \prec \bar{v}$. Therefore, $\bar{s} \prec \bar{\hat{s}} \prec \bar{v} \prec \bar{\hat{v}}$. Let $D_v$ denote the set of all descendant vertices of edges $\bar{e}_v$. From Lemma C.14, $D_v \subset \overline{Y}(s)$. Now, from the definition of hairpin, each vertex $u \in V_b$ such that $\bar{u} \in D_v$ must appear in an inverted closed walk in $G_b$ starting from $s$. Since the bracket set of $\bar{e}_v$ in $G_U$ is empty, such an inverted closed walk must include another nested inverted closed walk inside it that starts from $v$ and contains $u$. This implies each vertex in $Y(v)$ appears in an inverted closed walk starting from $v$. Furthermore, removing edge $e_v$ would disconnect $Y(v) \setminus \{v\}$ from the remaining graph. This means vertex $v$ also satisfies the hairpin conditions. But then $\langle s \rangle$ cannot be a hairpin by definition. We arrive at a contradiction.

**Theorem 1** *A vertex pair* $s, t \in V_b$ *satisfies all the panbubble conditions (excluding minimality) if and only if all the following conditions hold:*

- *P1.* *Either* $\bar{s} \prec \bar{\hat{s}} \prec \bar{\hat{t}} \prec \bar{t}$ *or* $\bar{t} \prec \bar{\hat{t}} \prec \bar{\hat{s}} \prec \bar{s}$ *in* $G_U$*.*
- *P2.* *Edges* $\bar{e}_s$ *and* $\bar{e}_t$ *are cycle equivalent in* $G_U$*.*
- *P3.* *Consider walks in* $G_b$ *that end at* $s$ *or* $t$ *but avoid passing through* $s$ *or* $t$*. Then, for every vertex* $u \in U(s,t) \cup U(t,s)$*, there exist such walks from* $u$ *to* $s$ *and from* $\hat{u}$ *to* $t$*, or there exist such walks from* $\hat{u}$ *to* $s$ *and from* $u$ *to* $t$*.*
- *P4.* *For every vertex* $v \in B(s,t) \setminus \{\hat{s}, \hat{t}\}$*, if edge* $\bar{e}_v$ *has an empty bracket set in* $G_U$*, then either* $\bar{e}_s \prec \bar{e}_v \prec \bar{e}_t$ *or* $\bar{e}_t \prec \bar{e}_v \prec \bar{e}_s$*.*

*Proof.* [$\Rightarrow$] If $\langle s, t \rangle$ is a panbubble in $G_b$, then conditions P1, P2, P3, and P4 are directly ensured by Lemma 2, Lemma 3, Lemma 4, and Lemma 5 respectively.

[$\Leftarrow$] For convenience, let us call a walk *special* if it ends at $s$ or $t$ and does not pass through $s$ or $t$. From the first condition P1, assume wlog that $\bar{s} \prec \bar{\hat{s}} \prec \bar{\hat{t}} \prec \bar{t}$ in $G_U$. Condition P3 implies that every vertex $u \in U(s,t) \cup U(t,s)$ appears in some special walk from $s$ to $t$. Thus, the traversability condition is satisfied. Condition P3 also implies that $U(s,t) = U(t,s)$. Therefore, the matching condition is also satisfied.

Next, we prove that the vertex pair $s, t$ satisfies the separable condition. Assume for contradiction that there exist vertices $v_{in} \in B(s,t)$ and $v_{out} \in V_b \setminus B(s,t)$ that remain connected after removing $e_s$ and $e_t$. If $v_{in}$ and $v_{out}$ are connected by a black edge, then both vertices would lie entirely within $B(s,t)$ or entirely outside it. Therefore, $v_{in}$ and $v_{out}$ must be connected by gray edge only.

From condition P3, suppose wlog that there exist special walks from $\hat{v}_{in}$ to $s$ and from $v_{in}$ to $t$. Vertex $v_{out}$ must be either $s$ or $t$; if not, there would exist special walks from $\hat{v}_{out}$ to $t$ passing through $v_{out}$ and $v_{in}$, which would imply

$v_{out} \in B(s,t)$. Therefore, $v_{out} \in \{s,t\}$. Now let us consider $G_U$. In $G_U$, (i) there exists a path starting from vertex $\bar{s}$ and edge $\bar{e}_s$ to $\bar{v}_{in}$ in $G_U$ without passing through $\bar{t}$, and (ii) there exists a path starting from vertex $\bar{t}$ and edge $\bar{e}_t$ to $\bar{v}_{in}$ in $G_U$ without passing through $\bar{s}$. An edge from $\bar{v}_{in}$ to either $\bar{s}$ or $\bar{t}$ would break the cycle-equivalence of edges $\bar{e}_s$ and $\bar{e}_t$, violating condition P2.

So far, we have established that the vertex pair $s, t$ satisfies the matching, traversability, and the separable condition. Finally, we show that the vertex pair $s, t$ satisfies the no-hairpin condition. Assume, for contradiction, that there exists a vertex $v \in B(s,t) \setminus \{\hat{s}, \hat{t}\}$ that serves as an entrance vertex of a hairpin. By Corollary C.8, it follows that $\bar{v} \prec \bar{\hat{v}}$ in $G_U$. From Lemma C.12, the bracket set of edge $\bar{e}_v$ is empty. Since our proof of Lemma C.10 depends solely on the matching, traversability, and separable conditions, we have $\bar{v} \in D_s \setminus D_t$, where $D_s$ and $D_t$ denote the set of all descendant vertices of $\bar{e}_s$ and $\bar{e}_t$, respectively. This observation, together with our prior assumptions and condition P4, imply that the ordering $\bar{e}_s \prec \bar{e}_v \prec \bar{e}_t$ must hold. This means $\bar{t} \in D_v \setminus \{\bar{\hat{v}}\}$, where $D_v$ is the set of all descendant vertices of edge $\bar{e}_v$, and hence $t \in Y(v) \setminus \{v, \hat{v}\}$ (by Lemma C.14). From Lemma C.15, the bracket set of edge $\bar{e}_t$ is non-empty and we also concluded that the bracket set of edge $\bar{e}_v$ is empty. Observe that this contradicts the cycle-equivalence of edges $\bar{e}_s$ and $\bar{e}_t$, i.e., condition P2.

**Proof of Theorem 2**

We state one auxiliary result before stating the proof of Theorem 2.

**Lemma C.16.** *Suppose $\langle s \rangle$ is a hairpin in $G_b$. Consider walks in $G_b$ that end at $s$ and avoid passing through $s$. For all vertices $u \in Y(s)$, there exist such walks originating from both $u$ and $\hat{u}$.*

*Proof.* We follow the same proof technique used for panbubbles in Lemma 4. Because $\hat{s} \in Y(s)$ appears on an inverted closed walk starting from $s$, the claim holds trivially when $u \in \{s, \hat{s}\}$. If $Y(s) \setminus \{s, \hat{s}\}$ is empty, we are done.

Suppose $Y(s) \setminus \{s, \hat{s}\} \neq \phi$, and consider a vertex $u \in Y(s) \setminus \{s, \hat{s}\}$. For convenience, let us call a walk *special* if it ends at $s$ and does not pass through $s$. By definition, vertex $u$ is reachable from $s$ without passing through $s$. Accordingly, there exist special walks from $u$ or $\hat{u}$ to $s$. If there exist special walks from both $u$ and $\hat{u}$ to $s$, then we are done. The other possibility is that there exist special walks from $u$ but not from $\hat{u}$ (or vice-versa). We will argue that this is not possible.

Assume for contradiction that special walks from $u$ exist, but no special walk from $\hat{u}$ exists. Since $\langle s \rangle$ is a hairpin and vertex $u \in Y(s) \setminus \{s, \hat{s}\}$, $u$ lies on an inverted closed walk $\omega$ starting from $s$. We consider two cases depending on whether $\hat{u}$ or $u$ appears first in $\omega$.

Case 1 (the first occurrence of $\hat{u}$ appears before the first occurrence of $u$ in $\omega$): Let us consider the sub-walk $\omega'$ of $\omega$ from the the first occurrence of $\hat{u}$ to the last vertex of $\omega$, i.e., $s$. From our earlier conclusion, there is no special walk from $\hat{u}$. Therefore, $\omega'$ must pass through $s$. Consider the first pass of this walk through black edge $e_s$. The corresponding ordered pair of vertices that appear in $\omega'$ must be one of $(s, \hat{s})$ or $(\hat{s}, s)$. Among these pairs, observe that pair $(\hat{s}, s)$ cannot occur because it would imply the existence of a special walk from $\hat{u}$. On the other hand, if pair $(s, \hat{s})$ occurs, then this would lead to violation of the separable property of hairpin $\langle s \rangle$. This leads to a contradiction, which confirms existence of special walks from both $u$ and $\hat{u}$ to $s$.

Case 2 (the first occurrence of $\hat{u}$ appears after the first occurrence of $u$ in $\omega$): Recall that walks in a biedged graph are symmetric, i.e., any walk from one vertex to another can be reversed to obtain a walk in the opposite direction. Let us consider the sub-walk $\omega'$ of $\omega$ from the the first occurrence of $\hat{u}$ in $\omega$ to the start of $\omega$, i.e., vertex $s$. One can now repeat the same arguments as Case 1 to arrive at a contradiction.

**Theorem 2** *A vertex $s$ is an entrance vertex of a hairpin in $G_b$ if and only if all the following hold:*

- *H1. The bracket set of edge $\bar{e}_s$ is empty in $G_U$.*
- *H2. There does not exist any vertex $v \in V_b \setminus \{\hat{s}\}$ such that $\bar{s} \prec \bar{v}$ in $G_U$ and the bracket set of edge $\bar{e}_v$ is empty.*
- *H3. Consider walks in $G_b$ that end at $s$ and avoid passing through $s$. For all vertices $u \in Y(s)$, there exist such walks from both $u$ and $\hat{u}$.*

*Proof.* [$\Rightarrow$] If vertex $s \in V_b$ is an entrance vertex of a hairpin in $G_b$, then condition H1 holds due to Lemma C.12. Condition H2 holds due to Lemmas C.14 and C.15. Finally, condition H3 holds due to Lemma C.16.

[$\Leftarrow$] For convenience, let us call a walk *special* if it ends at $s$ and does not pass through $s$. From condition H3, observe that there exist special walks from $s$ to $s$ in which vertex $u$ appears for all $u \in Y(s)$. In other words, $\forall u \in Y(s)$, $u$ belongs to an inverted closed walk starting from $s$. Hence, the first condition in the hairpin definition is satisfied. Next, we will argue that the second separable condition holds.

Assume for contradiction that there exist vertices $v_{in} \in Y(s) \setminus \{s\}$ and $v_{out} \in V_b \setminus (Y(s) \setminus \{s\})$ which remain connected after removing edge $e_s$. If $v_{in}$ and $v_{out}$ are connected by a black edge, then both vertices must entirely belong to one partition, either within $Y(s) \setminus \{s\}$ or its complement. Therefore, $v_{in}$ and $v_{out}$ must be connected by gray edge only.

From condition H3, there exist special walks from both $\hat{v}_{in}$ and $v_{in}$. Vertex $v_{out}$ must be equal to $s$; if not, there would exist special walks from $\hat{v}_{out}$ passing through $v_{out}$ and $v_{in}$, which would imply that $v_{out} \in Y(s) \setminus \{s\}$. Therefore, $v_{out} = s$. As there exist special walks from $s$ to $s$ in which $v_{in}$ appears, there also exist paths starting from vertex $\bar{s}$ and edge $\bar{e}_s$ to $\bar{v}_{in}$ in $G_U$ without passing through $\bar{s}$. An edge between $\bar{v}_{in}$ and $\bar{v}_{out} = \bar{s}$ would imply that vertex $\bar{s}$ is part of a cycle in $G_U$ that includes vertex $\bar{v}_{in}$. However, this is not possible because condition H1 ensures that the bracket set of edge $\bar{e}_s$ in $G_U$ is empty.

So far, we have established that vertex $s$ satisfies the first two conditions of the hairpin definition. To complete the proof, we need to argue that no vertex in $Y(s)$ other than $s$ satisfies the first two conditions from the hairpin definition. Assume, for contradiction, that there exists a vertex $v \in Y(s) \setminus \{s\}$ that satisfies the conditions. Because our proofs of Corollary C.9 and Lemma C.12 relied only on the first two conditions from the hairpin definition, we conclude that $\bar{s} \prec \bar{v}$ in $G_U$, and the bracket set of edge $\bar{e}_v$ is empty. But this contradicts H2. Hence, $s$ is the entrance vertex of a hairpin.

#### Proof of Theorem 3

**Theorem 3** *Let $\langle x, y\rangle$ and $\langle m, n\rangle$ be two different panbubbles such that $B(x,y) \cap B(m,n) \neq \phi$, then either $B(x,y)$ is a strict subset of $B(m,n)$ or $B(m,n)$ is a strict subset of $B(x,y)$.*

*Proof.* Since $\langle x, y\rangle$ and $\langle m, n\rangle$ are two different panbubbles, $x \neq y \neq m \neq n$ (using Lemma C.2). Suppose wlog that $\bar{e}_x \prec \bar{e}_y$ and $\bar{e}_m \prec \bar{e}_n$ (and hence by Lemma C.5, we have $\bar{x} \prec \overline{\hat{x}} \prec \overline{\hat{y}} \prec \bar{y}$ and $\bar{m} \prec \overline{\hat{m}} \prec \overline{\hat{n}} \prec \bar{n}$, respectively). Consider any vertex $v$ in $B(x,y) \cap B(m,n)$. From Corollary C.6, it follows that $\bar{x} \prec \bar{v}$ and $\bar{m} \prec \bar{v}$. Therefore, $\bar{x}$ and $\bar{m}$ share a root-leaf path, which means either $\bar{x} \prec \bar{m}$ or $\bar{m} \prec \bar{x}$. Suppose wlog that $\bar{x} \prec \bar{m}$.

We will prove that $B(m,n) \subset B(x,y)$. Assume for contradiction that $B(m,n) \not\subset B(x,y)$. Since $\bar{x} \prec \bar{m}$, we have $\bar{x} \prec \overline{\hat{x}} \prec \bar{m} \prec \overline{\hat{m}}$ and $\bar{e}_x \prec \bar{e}_m$. Let us use $D_x, D_y$, and $D_m$ to denote the set of all descendant vertices of edges $\bar{e}_x$, $\bar{e}_y$, and $\bar{e}_m$, respectively. Clearly, $\bar{y}, \overline{\hat{y}} \in D_x$. Let us consider the different possible positions of vertex $\bar{y}$ relative to tree edge $\bar{e}_m$ in $T_{v_{src}}$ (also see Figure S6A). Either (i) $\bar{x} \prec \bar{y} \prec \bar{m}$, (ii) $\bar{y}$ and $\bar{m}$ do not share any root-to-leaf path, or (iii) $\bar{m} \prec \bar{y}$. Condition (i) cannot hold because $\bar{e}_x \prec \bar{e}_y \prec \bar{e}_m$ implies $B(x,y) \cap B(m,n) = \phi$ (Lemma C.10). If (ii) holds, then all vertices in set $\overline{B}(m,n)$ belong to $D_x \setminus D_y = \overline{B}(x,y)$ (Lemma C.10) and we are done. Henceforth, we will assume that (iii) holds, that is, $\bar{m} \prec \bar{y}$. If $\bar{m} \prec \bar{y}$ and $\bar{y} = \overline{\hat{m}}$, then again $B(x,y) \cap B(m,n) = \phi$ (Lemma C.10). Thus, we assume $\bar{y} \neq \overline{\hat{m}}$. Accordingly, $\bar{e}_x \prec \bar{e}_m \prec \bar{e}_y$.

Let us now consider the possibilities of the positioning of vertex $\bar{n}$ relative to tree edge $\bar{e}_y$ in $T_{v_{src}}$ (also see Figure S6B). Either (a) $\bar{m} \prec \bar{n} \prec \bar{y}$, (b) $\bar{n}$ and $\bar{y}$ do not share any root-to-leaf path, or (c) $\bar{y} \prec \bar{n}$. If (a) holds, then all vertices in set $\overline{B}(m,n)$ belong to $D_x \setminus D_y = \overline{B}(x,y)$ (Lemma C.10) and we are done. Next, we will argue that (b) cannot hold. Assume for contradiction that (b) holds. All vertices in $D_y$ belong to $D_m \setminus D_n = \overline{B}(m,n)$ (Lemma C.10). Observe that $\bar{e}_y$ must have an empty bracket set. This is because $\bar{e}_x$ and $\bar{e}_y$ have identical bracket sets (Lemma C.7), and having an edge between a descendant of $\bar{e}_y$ to an ancestor of $\bar{e}_x$ would contradict the separable condition of $\langle m, n\rangle$. Now, the traversability condition of $\langle m, n\rangle$ suggests that for all vertices $v \in V_b$ such that $\bar{v} \in D_y$, there exists a walk in $G_b$ from $m$ to $n$ passing through $v$. Since $\bar{e}_y$ has an empty bracket set in $G_U$, this is only possible if there exists a hairpin entrance at vertex $u \in V_b$ and $\bar{u} \in D_y$. However, this would contradict the no-hairpin condition of $\langle m, n\rangle$.

Henceforth, we assume that (c) holds. As a result, $\bar{e}_x \prec \bar{e}_m \prec \bar{e}_y \prec \bar{e}_n$ (Figure S6C). In the remaining part of this proof, we will argue that $\bar{e}_x \prec \bar{e}_m \prec \bar{e}_y \prec \bar{e}_n$ is also not possible by contradiction. We will show that $\langle m, y\rangle$ is a panbubble if $\bar{e}_x \prec \bar{e}_m \prec \bar{e}_y \prec \bar{e}_n$, which would then contradict the minimality condition of $\langle x, y\rangle$ and $\langle m, n\rangle$.

Note that $\bar{e}_x$ and $\bar{e}_y$ have identical bracket sets (Lemma C.7). Similarly, $\bar{e}_m$ and $\bar{e}_n$ have identical bracket sets. Since $\bar{e}_x \prec \bar{e}_m \prec \bar{e}_y \prec \bar{e}_n$, it follows that $\bar{e}_x, \bar{e}_m, \bar{e}_y$, and $\bar{e}_n$ have identical bracket sets. Therefore, $\bar{m}$ or $\bar{y}$ must appear in a walk from a vertex in $D_m \setminus D_y$ to a vertex not in $D_m \setminus D_y$. In the following, we show that the vertex pair $(m, y)$ in $G_b$ satisfies all the panbubble conditions.

- *matching:* Assume for contradiction that $U(m,y) \neq U(y,m)$. Suppose wlog that $U(m,y) \setminus U(y,m) \neq \phi$. Choose any vertex $v$ from $U(m,y) \setminus U(y,m)$. Note that $v$ is reachable from $m$ without passing through $m$ or $y$. However, $v$ is not reachable from $y$ without passing through $m$ or $y$. As vertices $y$ and $\hat{y}$ belong to $U(y,m)$, $v$ is neither $y$ nor $\hat{y}$.

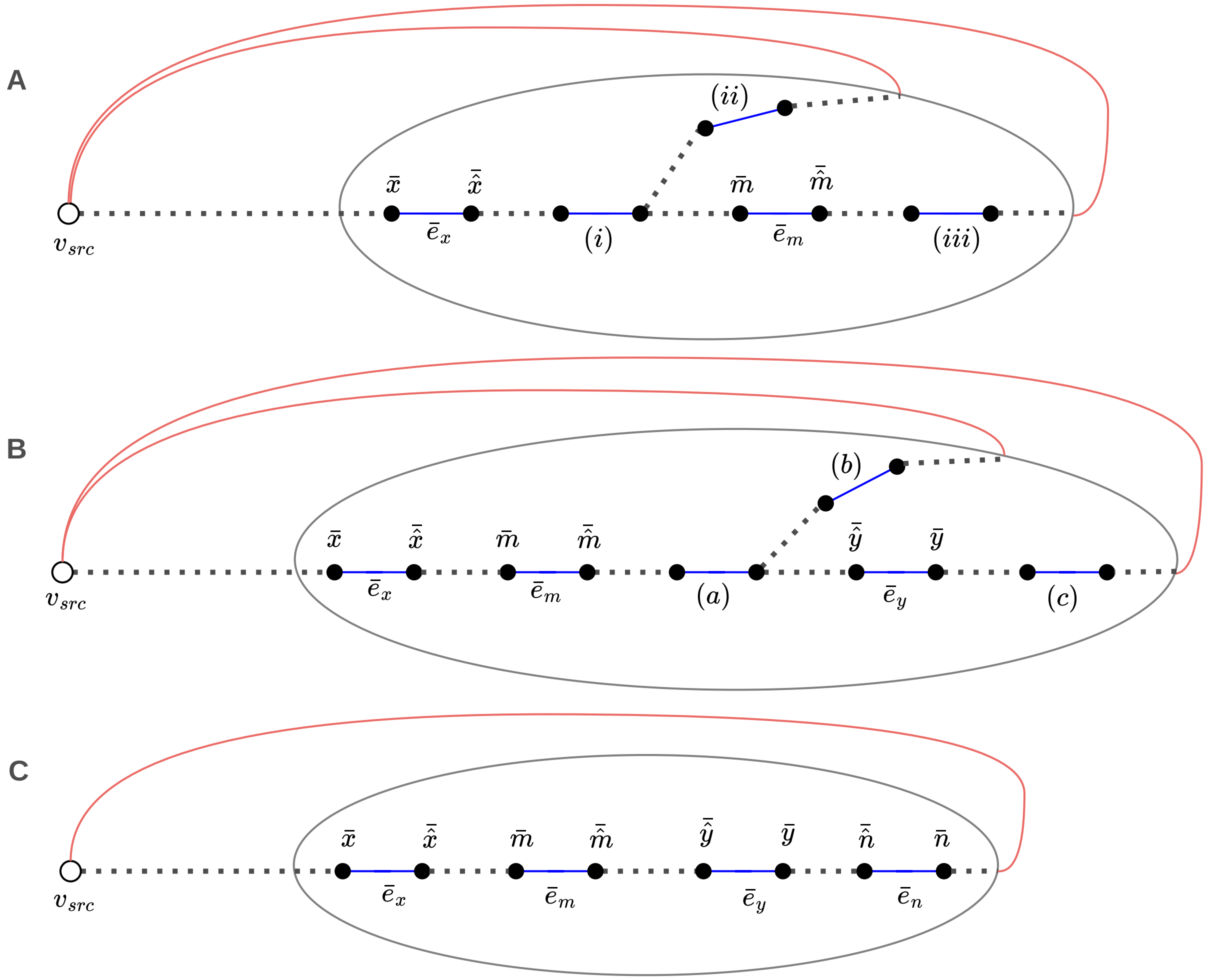


Fig. S6: Illustration of the case when $\bar{x} \prec \bar{m}$ in the proof of Theorem 3. **(A)** We depict the three possible positions of vertex $\bar{y}$ relative to tree edges $\bar{e}_x$ and $\bar{e}_m$. The three positions are labeled as (i), (ii), and (iii) in the figure. **(B)** We show the three possible positions of vertex $\bar{n}$ relative to tree edges $\bar{e}_m$ and $\bar{e}_y$. The three positions are labeled as (a), (b), and (c) in the figure. In **(C)**, we present the configuration satisfying $\bar{e}_x \prec \bar{e}_m \prec \bar{e}_y \prec \bar{e}_n$.

If $v$ is reachable from $m$ without passing through $m$ or $y$ in $G_b$, then $\bar{v}$ appears in some walk of type $(\bar{m}, \overline{\hat{m}}, \ldots)$ in $G_U$ which does not pass through $\bar{m}$ or $\bar{y}$. Since $\bar{e}_m \prec \bar{e}_y$, and $\bar{e}_m, \bar{e}_y$ have identical bracket sets, it follows that $\bar{v} \in D_m \cup \{\bar{m}\} \setminus D_y$.
From the above arguments, note that $v$ also belongs to $B(m, n)$ (Lemma C.10). If $\bar{e}_m, \bar{e}_y$ have identical bracket sets in $G_U$, then, in $G_b$, $v$ is reachable from $n$ without passing through $m$ or $n$ if only if $v$ is reachable from $y$ without passing through $m$ or $y$. Since $v \notin U(y, m)$, $v$ is not reachable from $n$ without passing through $m$ or $n$. This contradicts the matching condition of $\langle m, n \rangle$.

- *separable:* From the above arguments, it is clear that every vertex in $\overline{B}(m, y)$ belongs to the set $D_m \setminus D_y$. To prove separability, it suffices to show that $D_m \setminus D_y = \overline{B}(m, y)$. Assume for contradiction that there exists $v \in V_b$ such that $\bar{v} \in D_m \setminus D_y$ and $v$ is not reachable from $m$ without passing through $m$ or $y$. As $\bar{v} \in D_m \setminus D_y$, $v$ belongs to $B(x, y)$. Thus, $v$ is reachable from $x$ without passing through $x$ or $y$. In other words, there exists a walk from $x$ to $v$ or $\hat{v}$ without passing through $x$ or $y$. Since $\bar{e}_x, \bar{e}_m$ and $\bar{e}_y$ have identical bracket sets in $G_U$, this walk must pass through $m$ followed by $\hat{m}$ in $G_b$. This makes $v$ reachable from $m$ without passing through $m$ or $y$, leading to a contradiction.
- *traversability:* The above arguments also imply that $\overline{U}(m, y) \subset \overline{U}(x, y)$. Now since $\bar{e}_x, \bar{e}_m$ and $\bar{e}_y$ have identical bracket sets, all walks of type $(\bar{x}, \overline{\hat{x}}, \ldots \overline{\hat{y}}, \bar{y})$ in $G_U$ must include $\bar{m}$ immediately followed by $\overline{\hat{m}}$. In each of these walks, observe that the vertices that are in $D_m \setminus D_y$ must appear after the first appearance of $\bar{m}$. Recall that the traversability condition of $\langle x, y \rangle$ guarantees a walk in $G_b$ from $x$ to $y$ passing through $v$ for all $v$ in $U(x, y)$ (and hence for all $v$ in $U(m, y)$). It further guarantees a walk from $m$ to $y$ passing through $v$ for all $v \in U(m, y)$.
- *no-hairpin:* Since $U(m, y) \subset U(x, y)$, the presence of a hairpin at any vertex $v$ in $B(m, y) \setminus \{\hat{m}, \hat{y}\}$ would violate the no-hairpin condition of $\langle x, y \rangle$.
- *minimality:* If there is any vertex $v \in U(m, y)$ other than $y$ which pairs with $m$ and satisfies the above criteria, it would contradict the minimality condition of $\langle m, n \rangle$. Similarly, if there is any vertex $v \in U(m, y)$ other than $m$ which pairs with $y$ and satisfies the above criteria, it would contradict the minimality condition of $\langle x, y \rangle$.

Therefore, $\langle m, y \rangle$ is a panbubble. However, this contradicts the minimality condition of $\langle x, y \rangle$ and $\langle m, n \rangle$.

### Proofs of Theorem 4 and Theorem 5

The following claim directly holds due to the separable condition in our panbubble definition.

**Lemma C.17.** *Suppose $\langle s, t \rangle$ is a panbubble in biedged graph $G_b$. There is no gray edge in $G_b$ that connects vertices $s$ and $\hat{s}$, and there is no gray edge in $G_b$ that connects vertices $t$ and $\hat{t}$.*

**Theorem 4** *Let $\langle x, y \rangle$ and $\langle z \rangle$ be a panbubble and a hairpin respectively in $G_b$ such that $B(x, y) \cap (Y(z) \setminus \{z\}) \neq \phi$, then $B(x, y)$ is a strict subset of $Y(z) \setminus \{z\}$.*

*Proof.* Suppose wlog that $\bar{e}_x \prec \bar{e}_y$ in $G_U$. Therefore, $\bar{x} \prec \overline{\hat{x}} \prec \overline{\hat{y}} \prec \bar{y}$ (Lemma C.5). From Corollary C.8, we also have $\bar{z} \prec \overline{\hat{z}}$. Let us use $D_x, D_y$, and $D_z$ to denote the sets of all descendant vertices of edges $\bar{e}_x$, $\bar{e}_y$, and $\bar{e}_z$, respectively. From Lemma C.10 and Lemma C.14, $D_x \setminus D_y = \overline{B}(x, y)$ and $D_z = \overline{Y}(z) \setminus \{\bar{z}\}$.

We consider all possible positions of edge $\bar{e}_z$ relative to edges $\bar{e}_x$ and $\bar{e}_y$: (i) $\bar{e}_x$ and $\bar{e}_z$ do not share a root-to-leaf path, (ii) $\bar{e}_z \prec \bar{e}_x$, (iii) $\bar{e}_z = \bar{e}_x$, (iv) $\bar{e}_x \prec \bar{e}_z \prec \bar{e}_y$, (v) $\bar{e}_x \prec \bar{e}_z$ with $\bar{e}_y$ and $\bar{e}_z$ not sharing a root-to-leaf path, (vi) $\bar{e}_z = \bar{e}_y$, and (vii) $\bar{e}_x \prec \bar{e}_y \prec \bar{e}_z$. Below, we will argue that all the cases except (ii) are not possible.

Since panbubble $\langle x, y \rangle$ and hairpin $\langle z \rangle$ share one or more vertices (i.e., $B(x, y) \cap (Y(z) \setminus \{z\}) \neq \phi$), it is easy to observe that cases (i), (vi), and (vii) are not possible. Next, we consider case (iii). Using Lemma C.17 and Lemma C.12, we conclude that edge $\bar{e}_x$ has an empty bracket set. This means $\bar{e}_y$ also has an empty bracket set (Lemma C.7). But this contradicts our earlier claim that the interior of a hairpin cannot contain an edge with an empty bracket set (Lemma C.15). Lastly, cases (iv) and (v) are not possible because they contradict the no-hairpin condition of $\langle x, y \rangle$.

We are only left with case (ii), in which $\bar{e}_z \prec \bar{e}_x \prec \bar{e}_y$. In this case, observe that $D_x \setminus D_y = \overline{B}(x, y)$ is a strict subset of $D_z = \overline{Y}(z) \setminus \{\bar{z}\}$. Hence, the claim follows.

**Theorem 5** *Let $\langle m \rangle$ and $\langle n \rangle$ be two distinct hairpins in $G_b$. Then, the two hairpins do not share any vertex, i.e., $(Y(m) \setminus \{m\}) \cap (Y(n) \setminus \{n\})$ is an empty set.*

*Proof.* From Corollary C.8, we have $\bar{m} \prec \overline{\hat{m}}$ and $\bar{n} \prec \overline{\hat{n}}$ in $G_U$. Since $\langle m \rangle \neq \langle n \rangle$, $m \neq n$ and $e_m \neq e_n$. Now, either $\bar{e}_m$ and $\bar{e}_n$ share a root-to-leaf path or they do not. In the second case, it is clear that $(\overline{Y}(m) \setminus \{\bar{m}\}) \cap (\overline{Y}(n) \setminus \{\bar{n}\})$ is an empty set (Lemma C.14) and we are done.

Now consider the first case where $\bar{e}_m$ and $\bar{e}_n$ share a root-to-leaf path. We argue that this case is not possible. Suppose wlog that $\bar{e}_m \prec \bar{e}_n$. From Lemma C.12, bracket set of edge $\bar{e}_n$ is empty. Using Lemma C.11, we know that $\bar{n} \in \overline{Y}(m) \setminus \{\bar{m}, \overline{\hat{m}}\}$. However, this contradicts our earlier claim that such a vertex cannot be present in the interior of a hairpin (Lemma C.15).

### D Algorithms to detect panbubbles and hairpins: full exposition

#### Proof of Theorem 6

**Theorem 6** *Given a compact biedged graph $G(V_b, E_b)$, the entrance vertices of all panbubbles and hairpins can be enumerated exactly in $O(|V_b|^2(|V_b| + |E_b|))$ time. Moreover, the corresponding heuristic algorithm for the same task uses $O(|V_b|(|V_b| + |E_b|))$ time.*

*Proof.* Let us first analyze the exact algorithm for panbubble detection. Clearly, the initial steps of computing $G_U$, $T_{v_{src}}$, $A_d$, $A_f$, $A_{bridge}$, and the cycle-equivalent classes $C_1, C_2, \ldots, C_k$ require $O(|V_b| + |E_b|)$ time. Let $b_i$ denote the count of those edges in cycle-equivalent class $C_i$ whose corresponding edge in $G_b$ is black. Every pair of edges in this set of $b_i$ edges forms a candidate for a panbubble entrance.

In general, for any vertex pair $(s, t)$, where $s, t \in V_b$, the computation of arrays $W_s, W_t$ can be done in $O(|V_b| + |E_b|)$ time. By using these two arrays and our precomputed arrays $A_d$, $A_f$, and $A_{bridge}$, all the checks mentioned in Section 4.1 can be made in $O(1)$ time per vertex for all $v \in V_b$.

As a result, the total worst-case runtime to process cycle-equivalent class $C_i$ is $O(b_i^2(|V_b| + |E_b|))$. Thus, the time to process all $k$ cycle-equivalent classes is $O(\sum_{i=1}^{k} b_i^2(|V_b| + |E_b|))$. Now recall that for every vertex in $G_b$ there is exactly one black edge. Therefore, $\sum_{i=1}^{k} b_i$ is bounded by $|V_b|$, which further implies that $\sum_{i=1}^{k} b_i^2$ is bounded by $|V_b|^2$. Accordingly, the runtime complexity of detecting all panbubbles can be expressed as $O(|V_b|^2(|V_b| + |E_b|))$. Using a similar argument, it follows that the worst-case time complexity of the heuristic algorithm for detecting panbubbles is $O(|V_b|(|V_b| + |E_b|))$.

Next, we analyze the runtime of the exact algorithm for detecting hairpins. The computation of array $W$ to verify a hairpin entrance vertex can be done in $O(|V_b| + |E_b|)$ time. Using array $W$ and the other precomputed arrays, all the checks mentioned in Section 4.2 can be made in $O(1)$ time per vertex for all $v \in V_b$.

Every vertex in $G_b$ is a candidate for a hairpin entrance, and its verification takes $O(|V_b| + |E_b|)$ time by the above arguments. Therefore, the worst-case time complexity of detecting all hairpins is $O(|V_b|(|V_b| + |E_b|))$.

Note that both our exact and heuristic implementations compute hairpins exactly; they differ only in the algorithm used for computing panbubbles.
